# Activating Ras-MAPK pathway variants drive hippocampal clonal competition in human epilepsy

**DOI:** 10.64898/2026.01.26.701822

**Authors:** Sattar Khoshkhoo, Mingyun Bae, Yilan Wang, Ashton Tillett, Rosita B. Ramirez, Benjamin Finander, Emily D. Egan, Linus Marx, Dipan Patel, Zinan Zhou, Yasmine Chahine, Brian Chhouk, Sofia M. Zoullas, Abbe Lai, Roland Coras, Franck Bielle, Vincent Navarro, Bertrand Mathon, Taufik A. Valiante, Homeira Moradi Chameh, Andrew F. Gao, Timo Krings, Samuel Gooley, Michael S. Hildebrand, Kristian Bulluss, Jonathan Clark, Andrew P. Morokoff, James A. King, Marian Todaro, Patrick Kwan, Terence J. O’Brien, Samuel F. Berkovic, Ingrid E. Scheffer, Piero Perucca, Emily Lapinskas, John D. Rolston, G. Rees Cosgrove, Rani A. Sarkis, Alissa M. D’Gama, Sanda Alexadrescu, Edward Yang, Annapurna Poduri, R Mark Richardson, E. Zeynep Erson-Omay, Nihal DeLanerolle, Dennis D. Spencer, Katherine S.-M. Brown, Michael B. Miller, Amy E. Roberts, Luana N. Santos, Maria I. Kontaridis, Christian G. Bien, Stephen C. Blacklow, Kristopher T. Kahle, Ingmar Blümcke, August Yue Huang, Eunjung Alice Lee, Christopher A Walsh

## Abstract

Mesial (*a.k.a.,* medial) temporal lobe epilepsy (MTLE) is the most common focal epilepsy^1,2^ and, in drug-resistant cases, is treated by surgical removal of the anterior temporal lobe, which often shows neuronal loss and gliosis consistent with hippocampal sclerosis (HS)^2^. MTLE with HS has minimal contribution from germline genetic variation^3^, and is associated with prior precipitating insults such as prolonged childhood seizures and head trauma^4–6^. Somatic variants in Ras-MAPK pathway genes were recently reported in a few MTLE surgical specimens^7,8^, but their prevalence, clinical relevance, and underlying biological mechanisms remain unknown. Targeted duplex sequencing of hippocampal DNA from 462 surgical resections revealed significant enrichment of deleterious somatic variants in MTLE versus controls, with >40% of MTLE specimens harboring activating Ras-MAPK variants in *PTPN11*, *NF1*, *BRAF*, *KRAS*, and twelve genes not previously associated with focal epilepsy. Eight Ras-MAPK genes showed positive clonal selection in MTLE. Increased somatic variant burden predicted worse surgical outcome. Somatic Ras-MAPK variants at ultra-low (<0.5%) variant allele fractions were associated with older seizure onset and HS pathology, supporting a late prenatal or postnatal origin. Ras-MAPK variants in MTLE were enriched in cells derived from hippocampal progenitors—neurons, astrocytes, oligodendrocytes—in line with the known neuronal hyperexcitability and seizures induced by Ras-MAPK overactivation^9,10^; in contrast, Alzheimer disease hippocampi exhibited microglial enrichment of Ras-MAPK variants, consistent with prior reports^11^. Single-nucleus RNA sequencing showed increased expression of Ras-MAPK genes in neurons and upregulation of pathways mediating neurogenesis and neural development in MTLE. Functional validation of novel, recurrent *PTPN11* variants confirmed gain-of-function, while cellular modeling in induced pluripotent stem cells demonstrated proliferative/survival advantages for mutant cells in mosaic culture. Overall, our data suggest that somatic Ras-MAPK variants and acquired risk factors may converge on clonal competition in the hippocampus to modulate epilepsy risk.

## Introduction

Mesial temporal lobe epilepsy (MTLE), characterized by seizures arising from the hippocampus and nearby regions^1,2^, is the most common focal epilepsy. While many patients with MTLE experience significant seizure reduction on medications alone, roughly one-third of patients have drug-resistant disease and suffer from significant neurologic and psychiatric morbidity^12–14^. Surgical removal of the affected anterior temporal lobe has been used for nearly a century to control seizures in drug-resistant MTLE and is used widely in clinical practice today^1^. The most common histopathologic finding in MTLE surgical resections, hippocampal sclerosis (HS), is characterized by neuronal loss and gliosis in the hippocampal subregions^2^, and has been historically associated with acquired risk factors such as childhood complex febrile seizures, head trauma, and cerebral infections^2,4,5^. While germline genetic variants’ role in MTLE with HS is minimal^3,15^, post-zygotic (i.e., somatic) variants have emerged as a major contributor to other focal epilepsies^7,16–23^. Activating somatic variants in cancer-associated pathways like PI3K-mTOR and Ras-MAPK are now widely accepted as the causal agent for focal epilepsies associated with malformations of cortical development^16–20,23,24^ and low-grade epilepsy-associated tumors (LEATs)^21,22^, but their contribution to HS in the absence of tumor is unknown.

Hippocampal somatic variants in Ras-MAPK pathway genes were recently reported in a handful of cases of MTLE with HS^7,8,23,25^, but mainly in conjunction with temporal lobe focal cortical dysplasia (FCD) or LEAT dual pathology. Given the low variant allele fraction (VAF) of somatic Ras-MAPK variants in the absence of dysplasia or neoplasia (typically <2%)^7,8,25^, and limited sensitivity of conventional sequencing approaches at such low VAFs^26^, the burden and biological significance of these variants in MTLE with HS remain unknown. In contrast to FCDs that arise due to somatic mutation in early- to mid-gestation and typically cause epilepsy presenting in childhood^18,27^, MTLE is most common in adults and HS is strongly associated with precipitating risk factors^4,27^, implying a complex interaction between genetic and acquired factors in MTLE pathogenesis.

In this study, ultra-deep duplex sequencing with extensive experimental validation reveals a substantial burden of hippocampal deleterious Ras-MAPK pathway somatic variants in a large, international cohort of surgical MTLE tissue without dysplasia or neoplasia compared to age- and sex-matched neurotypical controls. These genetic findings correlate with key clinical and demographic information, including disease trajectory and outcome. Finally, single-nucleus RNA and cell type-specific DNA evaluation of human tissue, including comparison to Alzheimer’s disease (AD) samples, together with in vitro functional proteomics and induced pluripotent stem cell (iPSC) modeling experiments, suggest potential biological mechanisms of Ras-MAPK mosaicism in MTLE.

## Results

### Patient Cohort

Surgical samples from 462 patients with drug-resistant MTLE who underwent anterior temporal lobectomy for treatment of seizures were obtained from eight epilepsy surgical centers, following appropriate approvals by institutional ethics or review boards. The MTLE cohort included a range of characteristics representative of the clinical population (Supplementary Table 1), but cases with histopathological evidence of neoplasm in any part of the brain, including LEATs and FCD type IIIB, or dysplasia in the hippocampus were excluded. Post-mortem controls (Supplementary Table 1) were obtained from research brain banks and consisted of age- and sex-matched neurotypical individuals (Control, n=74, Extended Data Fig 1a-b) and AD patients with HS on histopathology (n=17).

### Targeted duplex sequencing of hippocampus-derived DNA

Hippocampus-derived DNA, extracted from fresh frozen brain tissue, underwent duplex sequencing using a custom panel of 71 genes, including genes involved in Ras-MAPK signaling^28,29^, genes associated with malformations of cortical development including FCD (PI3K-mTOR-FCD)^18–20^, and recurrent germline developmental and epileptic encephalopathy (DEE) genes^30^ (Extended Data Figure 1c). DNA libraries were sequenced, and raw reads sharing the same unique molecular identifiers (UMI) were aggregated to generate duplex reads (consensus between paired DNA strands), resulting in 1,836X and 1,206X deduplicated coverage, supported by at least one and two unique read pairs, respectively. MTLE, control, and AD samples showed comparable sequencing depths and most samples achieved >1,000X duplex coverage (Extended Data Fig 1d). Variant calling and pathogenicity enrichment were performed using pipelines benchmarked in prior studies^7,31^ (Fig 1a). For brevity, pathogenicity-enriched variants are referred to as “deleterious” in the text.

**Figure 1.**
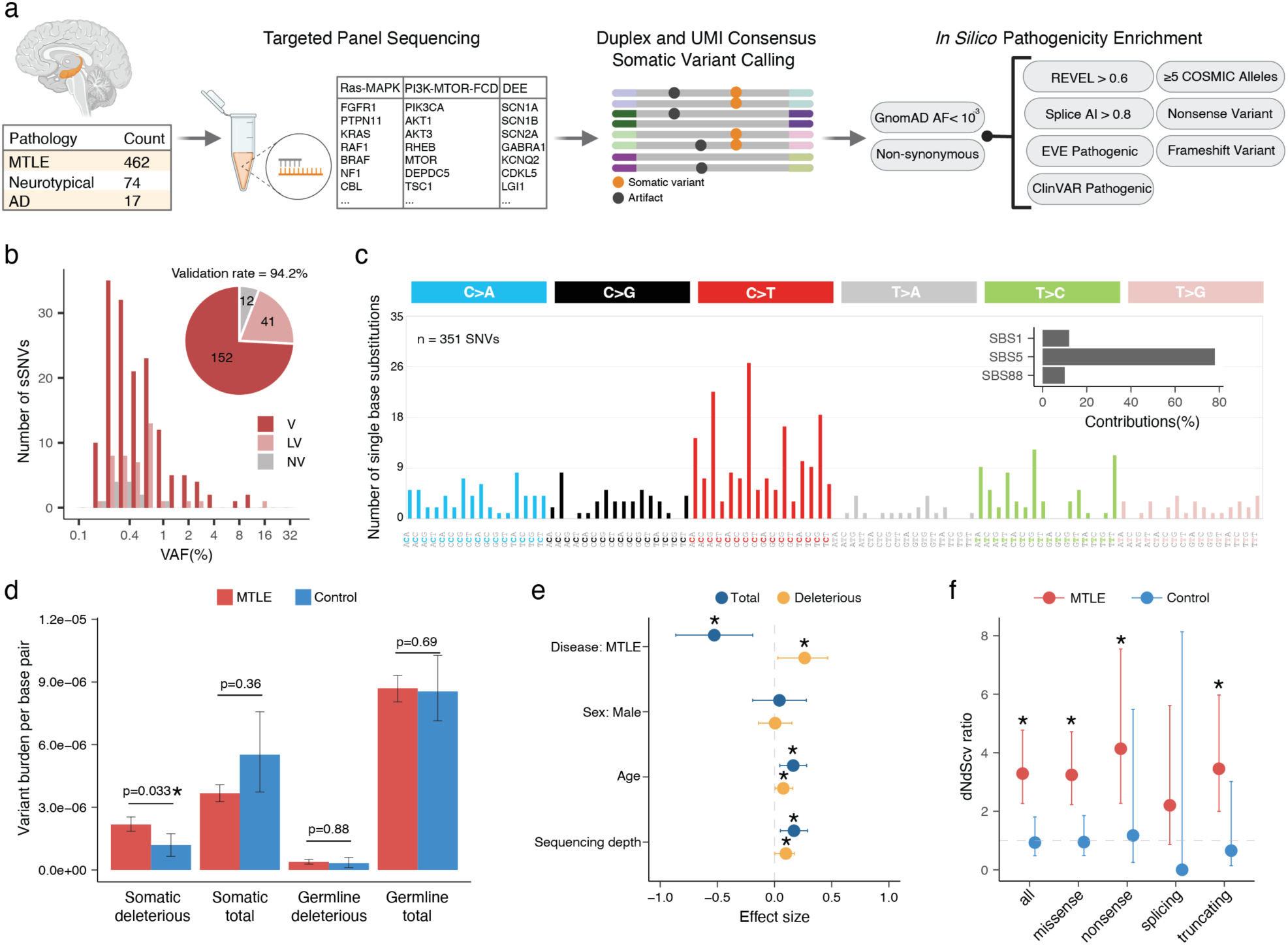
Overview of experimental approach and somatic variant burden in MTLE. (a) Study design: Duplex panel sequencing was used for high-fidelity identification of somatic variants in DNA derived from bulk hippocampal tissue from MTLE, AD, and neurotypical controls. *In silico* prediction tools were used to prioritize variants with increased likelihood of pathogenicity. (b) Validation rate and VAF distribution of orthogonally validated somatic variants. Variants are classified as validated (V), likely validated (LV), and not validated (NV) based on experimental validation. (c) Mutational spectrum of somatic single base substitutions across all samples. Top COSMIC mutational signatures are shown in the inset. (d) Somatic variant burden by variant type in MTLE and control samples (Two-sided Wilcoxon test; 95% confidence intervals [CI] via 1,000 bootstrap iterations). Asterisks indicate statistical significance. (e) Linear mixed-effects model analysis of somatic variant counts, showing contribution of disease status, sex, age, and sequencing coverage for all and deleterious somatic variants. Asterisks indicate statistical significance (95% CI non-overlapping with 0). (f) dN/dScv ratio across variant types. Dashed gray line represents neutral selection. Asterisks denote statistical significance (lower bound of the 95% CI > 1).

To maximize accuracy, a “stringent” callset of somatic single nucleotide variants (SNVs) and insertions-deletions (INDELs) was developed by requiring ≥2 unique duplex read pair support and applying other strict filters (Supplementary Table 2). 205 candidate somatic variants in the stringent callset were tested for validation using amplicon sequencing and/or droplet digital polymerase chain reaction (ddPCR). In total, 458 validation experiments were performed (Supplementary Table 3), and variants were categorized as validated (V), likely validated (LV), or not validated (NV) based on the strength of experimental evidence. Up to 94.2% of the stringent candidate somatic variants were orthogonally validated (V and LV; Supplementary Table 3, Fig 1b) with a strong correlation between the original sequencing and validation VAFs (Extended Data Fig 1e), confirming the accuracy of our variant calling and filtration strategy and its reliability for downstream analyses.

The trinucleotide context of SNVs showed enrichment of C>T substitutions, especially at CpG dinucleotides (Fig 1c), consistent with previous studies of developmental somatic variants in the brain^32^. Mutational signature analysis revealed greatest contributions from Catalogue Of Somatic Mutations in Cancer (COSMIC^33^) mutational signatures SBS1, SBS5, and SBS88 (Fig 1c, inset). SBS1 and SBS5 are associated with cell division and developmental timing respectively^34,35^ and SBS88 appears to be most active in the first decade of life^36^, overall suggesting hippocampal somatic variants are likely the result of both developmental and postnatal mutational processes.

### Enrichment and positive clonal selection of deleterious somatic Ras-MAPK variants in MTLE

While total somatic variant burden did not differ between MTLE and neurotypical control samples, deleterious somatic variant burden was significantly higher in MTLE (Fig 1d). Linear mixed-effect model testing the contribution of important covariates showed the greatest effect size for disease status (MTLE vs control), with small contributions from age and sequencing depth (Fig 1e). No differences in germline variant burdens were observed between the two cohorts (Fig 1d).

MTLE samples exhibited significantly increased non-synonymous to synonymous variant ratios (dN/dS ratio)—a marker of positive clonal selection in cancer^37,38^—overall and also specifically for missense, nonsense, and truncating variants (Fig 1f), whereas controls showed no evidence of clonal selection. Despite similar total somatic variant counts in the two cohorts, deleterious variants were enriched in MTLE, suggesting that they are under positive selection in the MTLE hippocampus, while functionally silent variants showed negative selection, perhaps reflecting the widespread neuronal cell death in HS.

Case-control comparison of somatic variants from the stringent callset revealed that, while deleterious somatic variants in Ras-MAPK genes were detected in some control brains, they were significantly enriched in MTLE (33% in MTLE vs 21% in control) and showed much stronger evidence of clonal selection in MTLE (Fig 2a-c, Extended Data Fig 2a-c). No significant case-control differences were observed in rates of PI3K-mTOR-FCD (4% vs 4%) or DEE (4% vs 3%) variants (Fig 2a-b, Extended Data Fig 2a). Gene-level analysis highlighted eight Ras-MAPK genes with significantly increased dN/dS ratios (q-value < 0.05) in MTLE that were nominally enriched in cases versus controls (*PTPN11, NF1, CBL, BRAF, RIT1, MAP2K1, KRAS, and SOS1*; Fig 2c). Combined together, significantly more deleterious variants were present in these genes in MTLE versus controls (OR = 2.1 and p = 0.018, Fisher’s exact test), although statistical significance was not achieved for individual genes given the smaller control cohort. In MTLE, oncogenes such as *PTPN11*, *RIT1, KRAS, MAP2K1*, and *BRAF* exhibited much higher missense- versus truncating-dN/dS ratios consistent with potential gain-of-function (GOF), while tumor suppressor genes such as *NF1*, *CBL,* and *LZTR1* exhibited higher truncating-dN/dS ratios supporting a likely loss-of-function (LOF) phenotype (Extended Data Fig 2d-f), overall indicating that most MTLE-associated deleterious somatic variants activate the Ras-MAPK pathway, which in turn drives clonal selection. Ras-MAPK genes with the greatest number of somatic variants—*PTPN11*, *NF1*, *CBL*, *BRAF*, and *RIT1*— exhibited multiple recurrent variants and clusters of variants in hotspots, providing additional evidence for pathogenicity (Extended Data Fig 2g-h). In fact, 127 MTLE-associated Ras-MAPK variants in the stringent dataset have already been reported as pathogenic or likely pathogenic in Clinical Variation Archive (ClinVar^39^; Supplementary Table 2), typically in the context of cancer or RASopathies, while their association with epilepsy is reported for the first time.

**Figure 2.**
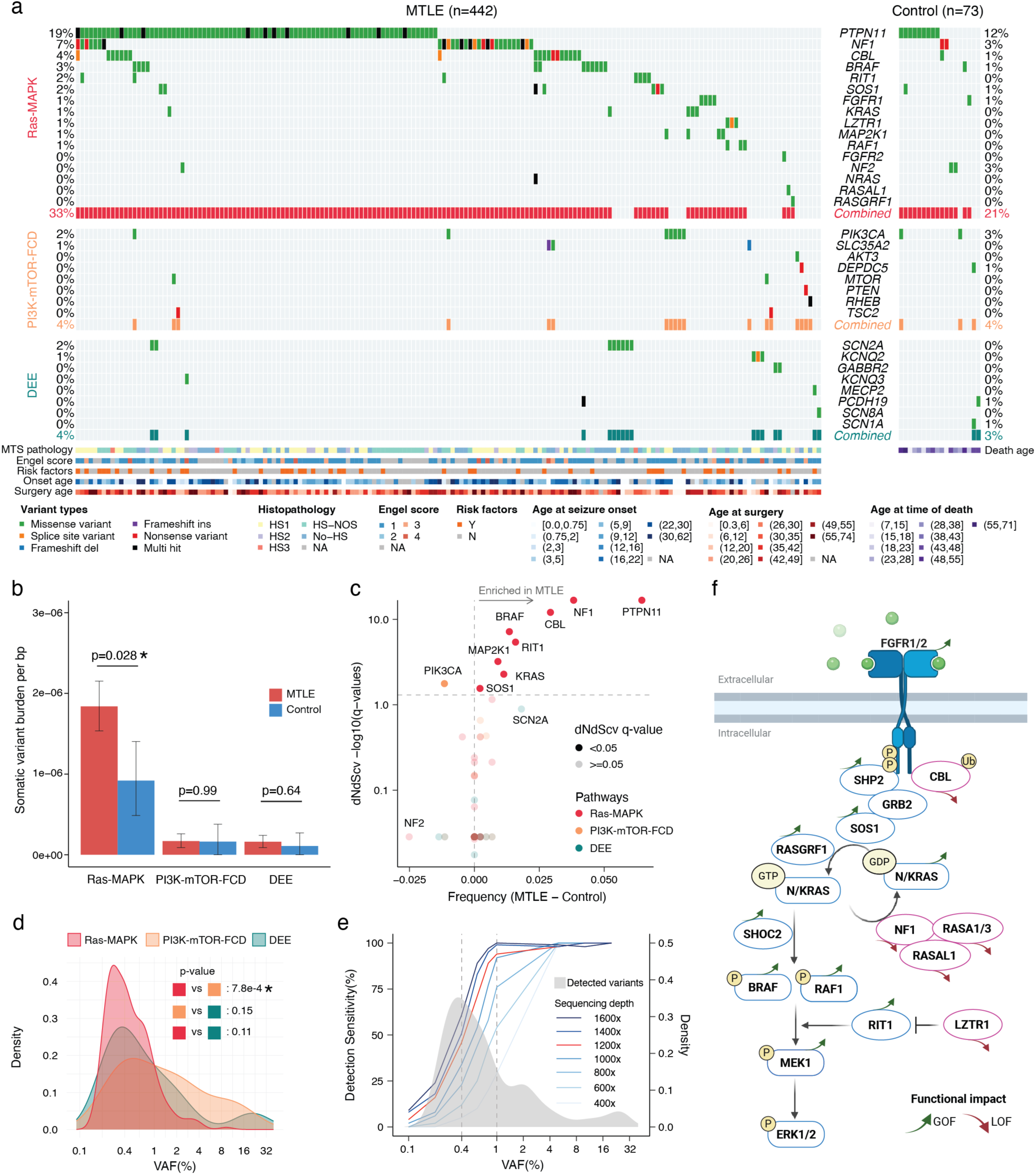
Spectrum of deleterious somatic variants in epilepsy-associated gene groups. (a) Oncoplot summarizing deleterious somatic variants detected in MTLE (n = 442) and controls (n = 73). Genes associated with Ras-MAPK signaling (top), PI3K-mTOR-FCD (middle), and DEE (bottom) are represented. The percentage of samples with deleterious somatic variants is shown for MTLE (left) and controls (right). Variant types and clinical metadata are shown at the bottom. (b) Variant burden per base pair by gene group (Two-sided Wilcoxon test; 95% CI via 1,000 bootstrap iterations). Asterisks indicate statistical significance. (c) Gene-level dN/dS analysis for MTLE variants. Genes to the right of the vertical dashed gray line are enriched in MTLE versus controls and genes above the horizontal dashed gray line show significantly increased nonsynonymous to synonymous variant ratios in MTLE (q-value < 0.05). (d) VAF distribution of deleterious somatic variants by gene group (Two-sided Wilcoxon test). Asterisks indicate statistical significance. (e) Simulation of somatic variant detection sensitivity by sequencing depth. Gray density plot shows VAF distribution of detected deleterious somatic variants. Simulated detection sensitivity by sequencing depth is color-coded in blue gradients, except for 1200X, which represents the average UMI>=2 sequencing depth in this study and is marked in red. (f) Simplified schematic of the Ras-MAPK signaling pathway. GOF and LOF MTLE variants in positive (blue) and negative (red) regulators of the pathway respectively, activate Ras-MAPK signaling.

### Low VAF and high prevalence of deleterious somatic Ras-MAPK variants in MTLE

Deleterious somatic Ras-MAPK variants in MTLE are present in a small fraction of cells (median VAF ∼0.4%), corresponding to small clones, with lower VAFs compared to PI3K-mTOR-FCD and DEE gene variants, suggesting a late gestational or an early postnatal origin (Fig 2d, Extended Data Fig 3a). Lower Ras-MAPK VAFs were not attributable to cohort-level differences, and sequencing depth did not account for the lower VAF or the higher abundance of Ras-MAPK variants (Extended Data Fig 3b-c). Low VAF variants were more frequently detected in regions with high sequencing depths suggesting potential underdetection of somatic variants in certain genomic loci despite high sequencing depths overall (Extended Data Fig 3d). Consistent with that, simulation of somatic variant detection sensitivity based on the stringent callset showed that only ∼50% of the somatic variants at the current median VAF of 0.4% are being detected (Fig 2e).

To increase somatic variant detection sensitivity, a second “sensitive” callset was generated (Supplementary Table 2) that also included: 1] variants with only one unique duplex read pair support, provided that for MTLE they were experimentally validated (Supplementary Table 3), 2] variants in samples with lower sequencing depths. Deleterious variants in the sensitive callset increased by ∼32% in MTLE (320 in the sensitive callset vs 243 in the stringent callset) and ∼23% in the controls (27 in the sensitive callset vs 22 in the stringent callset; Extended Data Fig 4a), with the fraction of MTLE samples carrying deleterious somatic Ras-MAPK variants increasing to 40.7% (versus 27.0% in the controls), while changes in the PI3K-mTOR-FCD and DEE gene variants were not significant (Extended Data Fig 4b-c). These values are still conservative since, unlike in the controls, candidate MTLE variants without experimental validation were not included in the sensitive callset. Genes with the largest increase in variant count in the sensitive callset—*PTPN11*, *NF1*, *CBL*, *BRAF*, and *SOS1* (Extended Data Fig 4d-e)—were among the most commonly mutated genes in MTLE, suggesting that somatic Ras-MAPK variants would be even more abundant in MTLE samples with deeper sequencing.

Our targeted duplex sequencing of MTLE hippocampi identified likely deleterious somatic variants in most canonical Ras-MAPK genes, predicted to increase pathway signaling through GOF of activators or LOF of repressors (Fig 2f). Somatic variants in genes such as *BRAF*, *FGFR1*, and *NF1*, have known associations with LEATs and large dysplasias^7,21–23^, but most Ras-MAPK variants and genes have never been described outside of tumor/dysplasia-related epilepsy. Specifically, we report 12 new candidate genes, such as *CBL*, *RIT1*, *LZTR1*, *RASA1*, *RASAL1*, and *RAF1*, along with >160 novel somatic Ras-MAPK variant associations with focal epilepsy.

### Association of somatic variants with MTLE risk factors and clinical findings

Retrospective histopathological evaluation of >200 cases and radiological evaluation of >100 cases showed that subjects with hippocampi harboring deleterious Ras-MAPK variants exhibited typical radiological and histopathological features associated with MTLE, mainly representing HS, without unique or aberrant features (Supplementary Table 1; some examples are shown in Extended Data Fig 5a-c). Notably, in a sample harboring the common BRAF p.V600E variant, immunostaining with a p.V600E-specific antibody confirmed the presence of this variant in neurons (Extended Data Fig 5a), and in a second case with the PTPN11 p.T52I variant, hypertrophic neurons were observed in the hippocampal cornu Ammonis (CA) 4 region with no other dysplastic features (Extended Data Fig 5c), consistent with prior reports^40,41^. Overall, MTLE hippocampi with histopathological evidence of HS had a similar burden of deleterious somatic variants to no-HS cases (mean [CI]: 0.58 [0.50, 0.60] and p = 0.33 in HS vs 0.48 [0.43, 0.68] and 0.29 in no-HS, two-sided Wilcoxon test; Fig 3a), but demonstrated significant enrichment of somatic Ras-MAPK variants and depletion of somatic PI3K-mTOR-FCD variants (odds ratio [OR] = 1.65 and p = 0.036 for Ras-MAPK, OR = 0.32 and p = 0.031 for PI3K-mTOR-FCD, Fisher’s exact test; Fig 3b). Among the MTLE risk factors, cases with a history of traumatic brain injury (TBI) had a higher, albeit non-significant, mean somatic variant burdens compared to cases without a history of TBI (0.92 [0.15, 1.08] and p = 0.10), but no clear trends were seen with febrile seizures (p = 0.99) or ischemic insults (p = 0.21, two-sided Wilcoxon test; Fig 3a). The presence of an MTLE risk factor was associated with lower likelihood of PI3K-mTOR-FCD mosaicism (OR = 0.21 and p = 0.024) without significant changes in other gene groups (p = 0.60 for Ras-MAPK and p = 0.62 for DEE, Fisher’s exact test; Fig 3b), indicating differential contribution of risk factors to clonal selection and evolution in the hippocampus. Cases with the worst epilepsy surgery outcome score, Engel score 4^42^, had the highest somatic variant burden (0.94 [0.22, 1.00] and p = 0.048, Fig 3a), without significant contribution from specific gene groups (p = 0.62 for Ras-MAPK, p = 0.53 for PI3K-mTOR, and p = 0.15 for DEE; Fig 3b), suggesting that higher deleterious somatic variant burden forecasts a lower likelihood of seizure-freedom after epilepsy surgery. These findings show that somatic variant-carrying hippocampi have a spectrum of typical histopathologic findings representing the norm and not unusual outliers, highlight potential causal relationships between somatic Ras-MAPK mosaicism and HS in the hippocampus, and suggest that possible contributions from MTLE risk factors such as TBI warrant further investigation.

**Figure 3.**
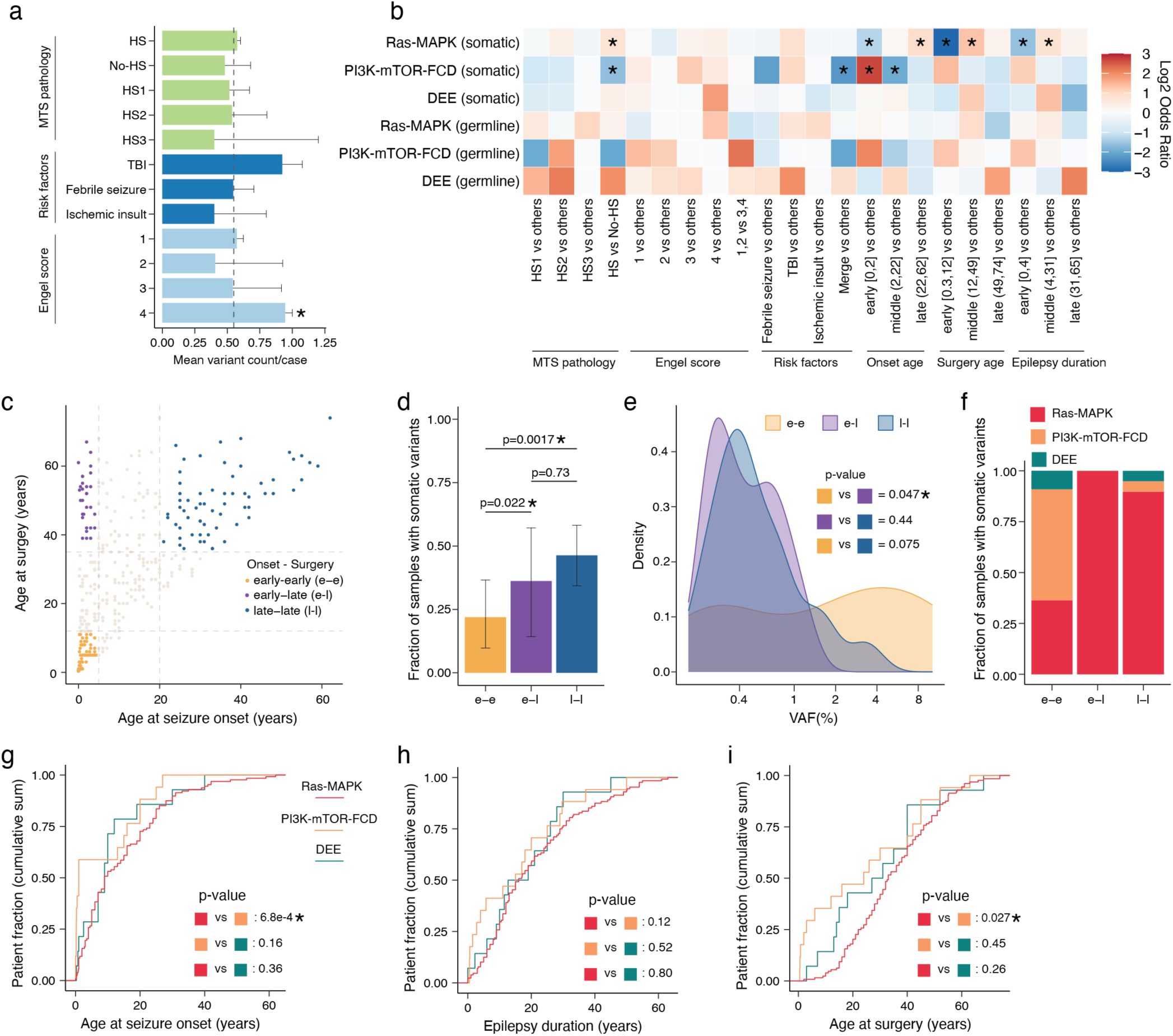
Association of deleterious somatic variants in MTLE with clinical and demographic variables. (a) Mean deleterious somatic variant count for specific clinical findings (Two-sided Wilcoxon test; 95% CI via 1,000 bootstrap iterations). Asterisks indicate p-values < 0.05. Dashed vertical line marks mean variant count for MTLE. (b) Association between somatic and germline variants and key clinical variables (Fisher’s exact test). Log2(odds ratio) is shown; asterisks indicate p-values < 0.05. (c) Distinct MTLE clinical subgroups based on age at seizure onset and age at surgery: e-e (n = 62), e–l (n = 30), and l–l (n = 64). (d) Proportion of samples with deleterious somatic variants in the e–e, e–l, and l–l groups (Two-sided Wilcoxon test, 95% CI via 1,000 bootstrap iterations). Asterisks indicate statistical significance. (e) VAF distribution of deleterious somatic variants in the e–e, e–l, and l–l groups (Two-sided Wilcoxon test). Asterisks indicate statistical significance. (f) Relative proportion of deleterious somatic variants in epilepsy-associated pathways for the e–e, e–l, and l–l groups. (g-i) Cumulative sum distributions of age at onset (g), epilepsy duration (h), and age at surgery (i; Kolmogorov–Smirnov test). Asterisks indicate statistical significance.

### Somatic Ras-MAPK variants correlate with later seizure onset and longer epilepsy duration

Analysis of age of seizure onset and age at surgery revealed specific enrichment of Ras-MAPK variants in cases with later onset of epilepsy, as adolescents or adults, and later age at epilepsy surgery (Fig 3b). Study subjects were divided into three groups (early [first two deciles], middle [middle six deciles], late [last two deciles]) based on age at seizure onset, age at surgery, and time from seizure onset to surgery (duration), and the frequency of deleterious somatic variants in each group was tested (Fig 3b). Somatic Ras-MAPK and PI3K-mTOR-FCD variants had opposing age-related trends: Ras-MAPK variants were less common in individuals with seizure onset in early childhood (“early” OR = 0.41 and p = 0.0044) but were enriched in individuals with their first known seizure as adults (“late” OR = 1.87 and p = 0.020, Fisher’s exact test), while conversely somatic PI3K-mTOR-FCD variants were strongly associated with early childhood seizures (“early” OR = 6.34 and p = 3.8e-4) but had significantly lower frequency with later onset of seizures (“middle” OR = 0.25 and p = 0.010, Fisher’s exact test). Younger age of epilepsy surgery and shorter epilepsy duration showed lower somatic Ras-MAPK variant burden (“early” OR = 0.13 and p = 9.7e-5 for surgery age, “early” OR = 0.33 and p = 4.2e-4 for epilepsy duration), while samples from epilepsy surgery in adolescence and early adulthood and individuals with longer epilepsy durations harbored more of these variants (“middle” OR = 2.45 and p = 4.8e-5 for surgery age, “middle” OR = 1.67 and p = 0.027 for epilepsy duration, Fisher’s exact test). Opposite trends for age at surgery and epilepsy duration were observed for PI3K-mTOR-FCD variants, but none of the values reached statistical significance.

To further examine how age at the time of first seizure and age at surgery relate to hippocampal somatic variants, we defined three subgroups (Fig 3c) representing distinct clinical entities: 1] early-early (e-e) group developed epilepsy early (<5) and had surgery early (<12), 2] early-late (e-l) group had early seizure onset (<5), but had surgery later in life (>35), 3] late-late (l-l) group developed epilepsy late (>20) and underwent surgery later in life (>35). As before (Fig 1e), older surgical age correlated with increased fraction of samples with deleterious somatic variants (Fig 3d), with a skew toward lower VAFs suggesting later developmental origin (Fig 3e). Most somatic PI3K-mTOR-FCD variants were found in the e-e cohort, whereas somatic Ras-MAPK variants were enriched in the e-l and l-l groups (Fig 3f). Across the whole cohort, somatic PI3K-mTOR-FCD variants were associated with earlier seizure onset compared to somatic Ras-MAPK variants, whereas incidence of somatic Ras-MAPK variants increased with age and was not related to epilepsy duration (Fig 3g-i, Extended Data Fig 5d-f). These findings are likely related to the observation that frequency of HS did not depend on age at seizure onset, but covaried with age at surgery and was maximal in the age group that exhibited most significant somatic Ras-MAPK mosaicism (Extended Data Fig 5g-h). These data support a potential causal link between HS and somatic Ras-MAPK variation and suggest that unlike somatic PI3K-mTOR-FCD variants that arise mid-gestation^20,43^, somatic Ras-MAPK variants in MTLE may arise, or become clonally amplified, postnatally.

### Minimal contribution of germline variants in drug-resistant MTLE

The overall prevalence of potentially deleterious germline variants in all cohorts was low (<10%) and comparable in MTLE cases and controls (Extended Data Fig 6a-c). No significant differences in deleterious germline variant burden or fraction of samples carrying deleterious germline variants were seen among the different gene groups (Extended Data Fig 6b-c). The fraction of samples with co-occurring germline and somatic variants was however significantly higher in MTLE, and specifically enriched in Ras-MAPK genes, suggesting that certain germline variants may contribute to differences in somatic variation (Extended Data Fig 6d-g). In some cases, for example in patients with deleterious germline *NF1* variants (Extended Data Fig 6g), this finding is consistent with Knudson’s two-hit model for pathogenicity in tumor suppressor genes^44^. While cancer-independent co-occurrence of germline and somatic *NF1* variants has been previously reported^45^, to our knowledge this is the first observation in the hippocampus or in focal epilepsy in the absence of tumor. Interestingly, three individuals in the MTLE cohort with clinical diagnoses of neurofibromatosis type 1 and pathogenic germline *NF1* variants (MTLE099, MTLE348, and MTLE371) have 6, 3, and 6 deleterious somatic *NF1* variants detected in their hippocampi, respectively, indicating an ongoing and rapid process of positive clonal selection in the hippocampus. The two-hit model is not applicable to most other MTLE-associated somatic variants that are in oncogenes such as *PTPN11*, and typically only require a single variant for pathogenicity; although even for oncogenes there is evidence of multiple somatic hits in the same sample (Extended Data Fig 6f-g) suggesting similar clonal selection processes that may be a key pathogenic mechanism in MTLE.

### Subtype-specific neuronal loss and upregulation of Ras-MAPK genes in MTLE

MTLE shows profound histopathological changes in the hippocampus^2^, but the potential impact of Ras-MAPK signaling on MTLE pathology, particularly HS, is unknown, as cellular and molecular changes have not been studied at the single-cell level before. We performed single-nucleus RNA sequencing (snRNA-seq) on frozen hippocampi from 6 MTLE (5 HS and 1 no-HS) and 2 neurotypical controls, and integrated with published data from 3 additional MTLE (all no-HS) cases^46^, overall producing 106,343 nuclei (26,252 HS, 70,919 no-HS, and 9,172 control) that passed QC (Fig 4a-b). Data integration and cluster identification using known cell type markers were performed with high concordance across cases, cohorts, and HS pathologies (Fig 4b, Extended Data Fig 7a-c). Cell type proportion analysis showed significant differences between cases and controls (Fig 4c) with consistent findings across cases with the same pathology (Extended Data Fig 7d). While all MTLE hippocampi showed depletion of interneurons, HS cases had more significant loss of excitatory pyramidal cells (PC) and granule cells (GC) compared to controls (Fig 4d-e). Additionally, hippocampi with HS exhibited an enrichment of inflammatory cells such as microglia (MG), T cells (TC), and macrophages (MP; Fig 4e).

**Figure 4.**
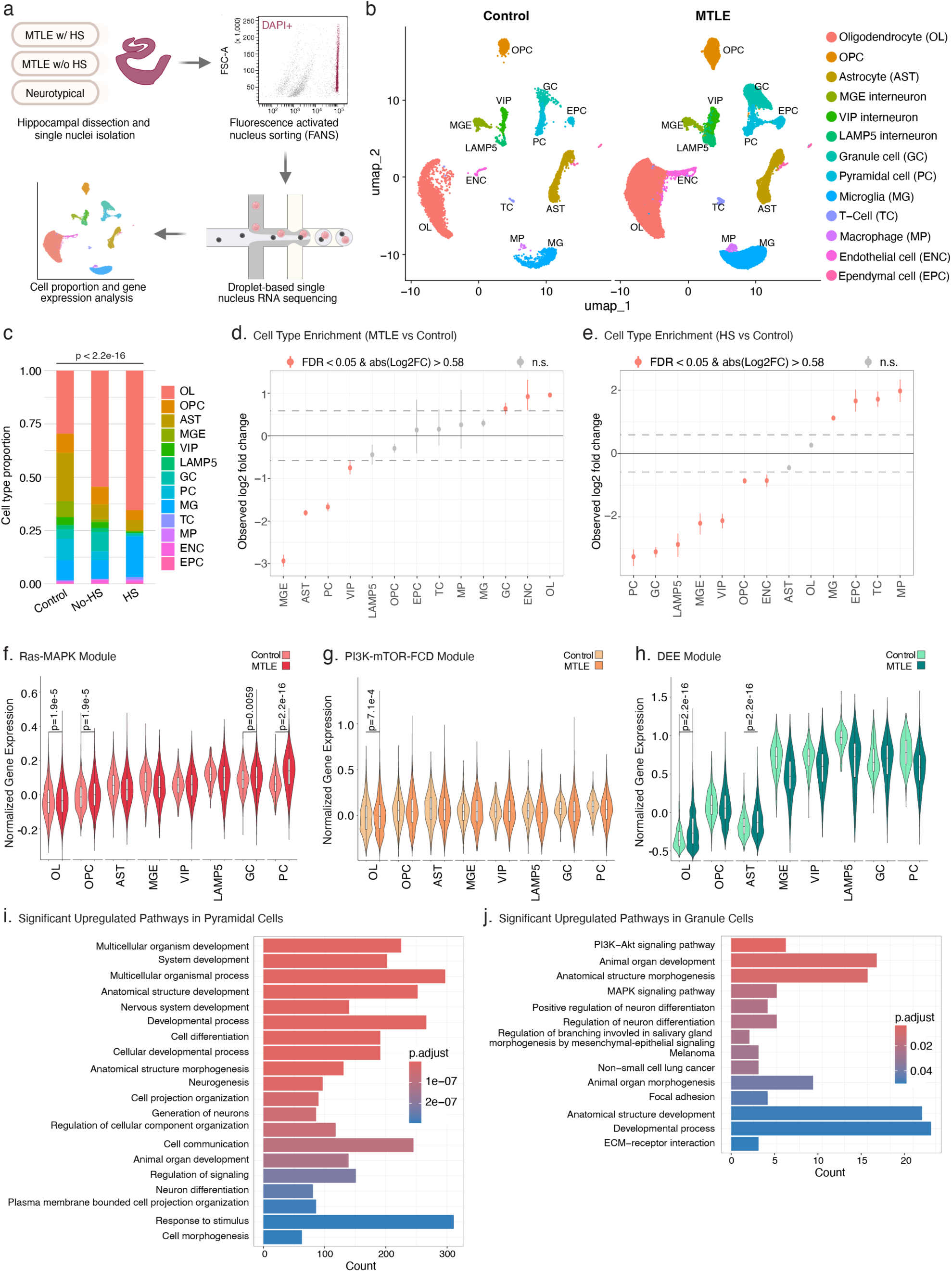
Cellular and transcriptional characteristics of MTLE. (a) snRNA-seq experimental workflow. (b) Unsupervised clustering of snRNA-seq data in control and MTLE hippocampi. (c) Cell type proportions based on snRNA-seq clusters in control and MTLE brains (with and without HS; Chi-squared proportion test for MTLE vs control). (d-e) Cell type enrichment/depletion analysis in all MTLE cases and in MTLE cases with HS vs controls (Two-sided Wilcoxon test for pairwise cell type proportion statistics; permutation test with1,000 permutations for fold difference statistics). Pink dots above and below the dashed lines represent enrichment or depletion in MTLE respectively. (f-h) Normalized expression levels for Ras-MAPK, PI3K-mTOR-FCD, and DEE genes in the targeted sequencing panel per cell type (Two-sided Wilcoxon test). Boxes represent median±25% quartiles; p-values <0.05 are shown. (i-j) Top differentially upregulated gene ontology pathways in MTLE vs controls for pyramidal and granule cells. Only gene pathways with Benjamini-Hochberg-adjusted p-values <0.05 are shown.

To confirm the potential functional impact of deleterious somatic variants, we tested expression levels of all the genes on the duplex sequencing panel in different cell types, showing that most panel genes were highly expressed in neurons (Fig 4f-h, Extended Data Fig 8a-f). However, case-control comparisons showed significantly higher expression of only Ras-MAPK genes, and not PI3K-mTOR-FCD or DEE genes, in excitatory neurons (Fig 4f-h). Further analysis showed differential expression of many genes in all the cell types (Supplementary Table 4; Extended Data Fig 8g), but gene ontology analysis showed the highest number of enriched pathways in excitatory neurons (Supplementary Table 5). In PC, pathways related to neurogenesis, neural development, and neuronal communication were most upregulated while ribosomal biogenesis and protein translation were most downregulated (Fig 4i, Extended Data Fig 8h). GC similarly showed upregulation of neural development and differentiation pathways including PI3K-Akt and MAPK signaling (Fig 4j), but most downregulated pathways were involved in ATP metabolism and oxidative phosphorylation (Extended Data Fig 8i). These findings overall highlight a key role for excitatory neurons in MTLE pathogenesis with most significant changes in gene programs regulating neurogenesis, and neural development and function, which may be impacted by Ras-MAPK signaling^47,48^.

### Specific enrichment of Ras-MAPK variants in hippocampal neural cell types in MTLE

Brain region- and cell type-specific genotyping of surgical tissue (Fig 5) shows that somatic Ras-MAPK variants in MTLE are present in cells derived from hippocampal neuronal progenitors. Analysis of DNA derived from the hippocampus and adjacent temporal neocortex in 17 Ras-MAPK variant-carrying MTLE cases, using ddPCR, showed that mutant alleles were limited to the hippocampus for 16/17, and enriched >10-fold in the hippocampus in the 17^th^ sample (Supplementary Table 6, Fig 5a), suggesting that 16/17 variants arose after separation of hippocampal and neocortical progenitor cells at mid-gestation. Florescence activated nucleus sorting (FANS) of surgical tissue paired with ddPCR genotyping (Fig 5b, Extended Data Fig 9a-b) showed that somatic Ras-MAPK variants were present in combinations of neurons, astrocytes, and oligodendrocytes, indicating origin in hippocampal neural progenitors—perhaps in the dentate gyrus (DG)—that produce these cell types (Fig 5b). Fewer neurons relative to glial cells harboring somatic variants is consistent with postnatal clonal expansion of glial cells and extensive hippocampal neuronal loss in HS. The specificity of neural enrichment of somatic Ras-MAPK variants in MTLE was confirmed by FANS-genotyping in microglia, that arise from a non-neural lineage (Extended Data Fig 9c-d). Consistent with prior reports^11^, AD samples with HS pathology showed enrichment of somatic Ras-MAPK variants in the microglia, while somatic variants were not detected in the microglia in MTLE (Extended Data Fig 9d). Cell type analysis of Ras-MAPK variants in control samples was limited by tissue availability and the very low VAF that precluded consistent detection in sorted cells. Selective enrichment of somatic variants in neural cell types in the hippocampus is congruent with existing models of epileptogenesis in MTLE^49–52^ and implicates the DG, which shows ongoing neurogenesis after birth in humans^53–62^, as a potential source of deleterious somatic variants with low VAF.

**Figure 5.**
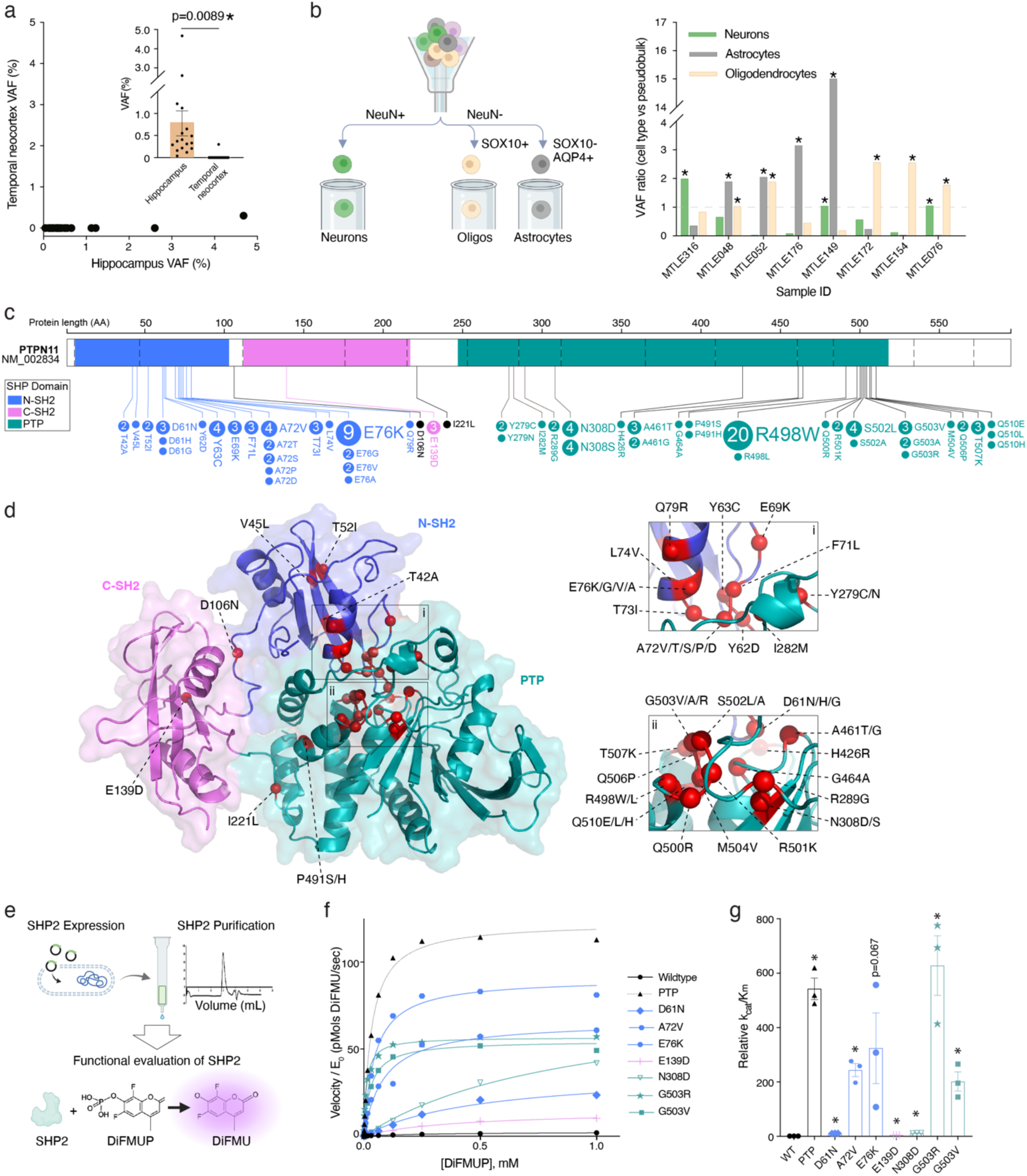
Cellular origins and functional characteristics of Ras-MAPK somatic variants in MTLE. (A) Enrichment of MTLE-associated deleterious somatic variants in the hippocampus relative to the temporal neocortex in the same individual. Inset shows % VAF distribution in different brain regions (Two-tailed unpaired t-test). Mean±SEM is shown. Asterisks indicate statistical significance. (B) Nuclear sorting strategy to isolate neurons, astrocytes, and oligodendrocytes in MTLE samples and the relative abundance of somatic variants in each neuroglial cell type as measured by ddPCR. Cell types with VAFs greater than the pseudobulk value (ratio of cell type-specific VAF over mean VAF for all cell types > 1, marked with asterisks) show relative enrichment in each sample. (c) Graphical representation of amino acid changes corresponding to MTLE-associated *PTPN11* variants along the gene body. Colors indicate SHP2 functional domains. Dashed vertical lines mark exon boundaries. Numbers inside circles show variant counts if >1. (d) Amino acids superimposed on the 3D structure of SHP2. Figure insets provide zoomed-in views of mutational hotspots in the catalytic domain of the enzyme. (e) Overview of DiFMUP assay for evaluation of SHP2 phosphatase activity. (f) Catalytic activity of SHP2^WT^, the isolated catalytic domain (PTP), and various SHP2^MUT^ as a function of substrate concentration (n = 3 replicates). Mean±SEM and non-linear best fit are shown; *R*^2^: WT (0.99), PTP (0.99), D61N (0.98), A72V (0.98), E76K (0.99), E139D (0.99), N308D (0.99), G503R (0.97), G503V (0.98). (g) Catalytic efficiency of each SHP2^MUT^ relative to SHP2^WT^ (n =3 replicates; two-tailed unpaired t-test for comparison against SHP2^WT^). P-values: PTP (1.6e-4), D61N (3.8e-6), A72V (5.4e-4), E76K (0.0674), E139D (9.1e-6), N308D (2.0e-6), G503R (0.0046), G503V (0.0046). Mean±SEM is shown; asterisks indicate p<0.05.

### MTLE-associated *PTPN11* variants increase SHP2 protein activity and drive positive clonal selection

*PTPN11* showed the most deleterious variants in MTLE with 111 variants resulting in 52 unique amino acid changes (Supplementary Table 2, Fig 5c), but many of these variants were novel with unknown functional significance. Analysis of the mutated amino acids in the SHP2 protein, which is encoded by *PTPN11*, showed clustering of most variants in two primary hotspots in the N-terminal Src homology-2 (N-SH2) and protein tyrosine phosphatase (PTP) domains of the protein (Fig 5c). Because most MTLE-associated variants map to this N-SH2-PTP interaction site (Fig 5d), they are predicted to disrupt the natural autoinhibited state of the protein, resulting in GOF^63^. This prediction was directly tested by comparing phosphatase activity of several recurrent SHP2 variants (SHP2^MUT^) against the wildtype protein (SHP2^WT^) and the isolated PTP domain *in vitro*. SHP2^MUT^, SHP2^WT^, and PTP were expressed, purified, and then incubated with a fluorogenic substrate DiFMUP (6,8-Difluoro-4-Methylumbelliferyl Phosphate) to quantify phosphatase activity (Fig 5e). All tested variants (D61N, A72V, E76K, E139D, N308D, G503R, G503V) exhibited increased basal phosphatase activity relative to SHP2^WT^ (Fig 5f). MTLE-associated SHP2^MUT^, irrespective of the affected protein domain, also resulted in increased catalytic efficiency (Fig 5g). These experimental results support a GOF phenotype for SHP2^MUT^, which is consistent with the broader assertion that somatic Ras-MAPK variants in MTLE result in overactivation of the pathway.

To test cell autonomous effects of Ras-MAPK somatic variants on clonal competition, mosaic induced pluripotent stem cell (iPSC) cultures were created by mixing an engineered PTPN11 p.G503R cell line and its isogenic wildtype control line at 0.1%, 5%, and 50% mosaicism (Fig 6a). Relative abundance of mutant and wildtype alleles was measured over time using ddPCR, providing an indirect measurement of cell fraction and clonal selection. At all starting mosaic ratios, the mutant cell fraction consistently increased over time, indicating increased clonal fitness and/or proliferation of the mutant cells. We then differentiated the PTPN11 p.G503R and isogenic wildtype iPSC lines into neural progenitor cells and facilitated spontaneous terminal differentiation into neural lineage cells. Consistent with the cell type-specific genotyping experiments in human MTLE tissue, the mutant cells gave rise to all major neuroglial cell types including neurons, astrocytes, and oligodendrocytes (Fig 6b). Given the association of MTLE-associated risk factors with aberrant DG neurogenesis in animal models^51,64–67^, these data provide a framework for how variant-carrying neural progenitors may outcompete the wildtype clones in the hippocampus and produce a disproportionately higher number of mutant neurons and glial cells that ultimately contribute to epileptogenesis.

**Figure 6.**
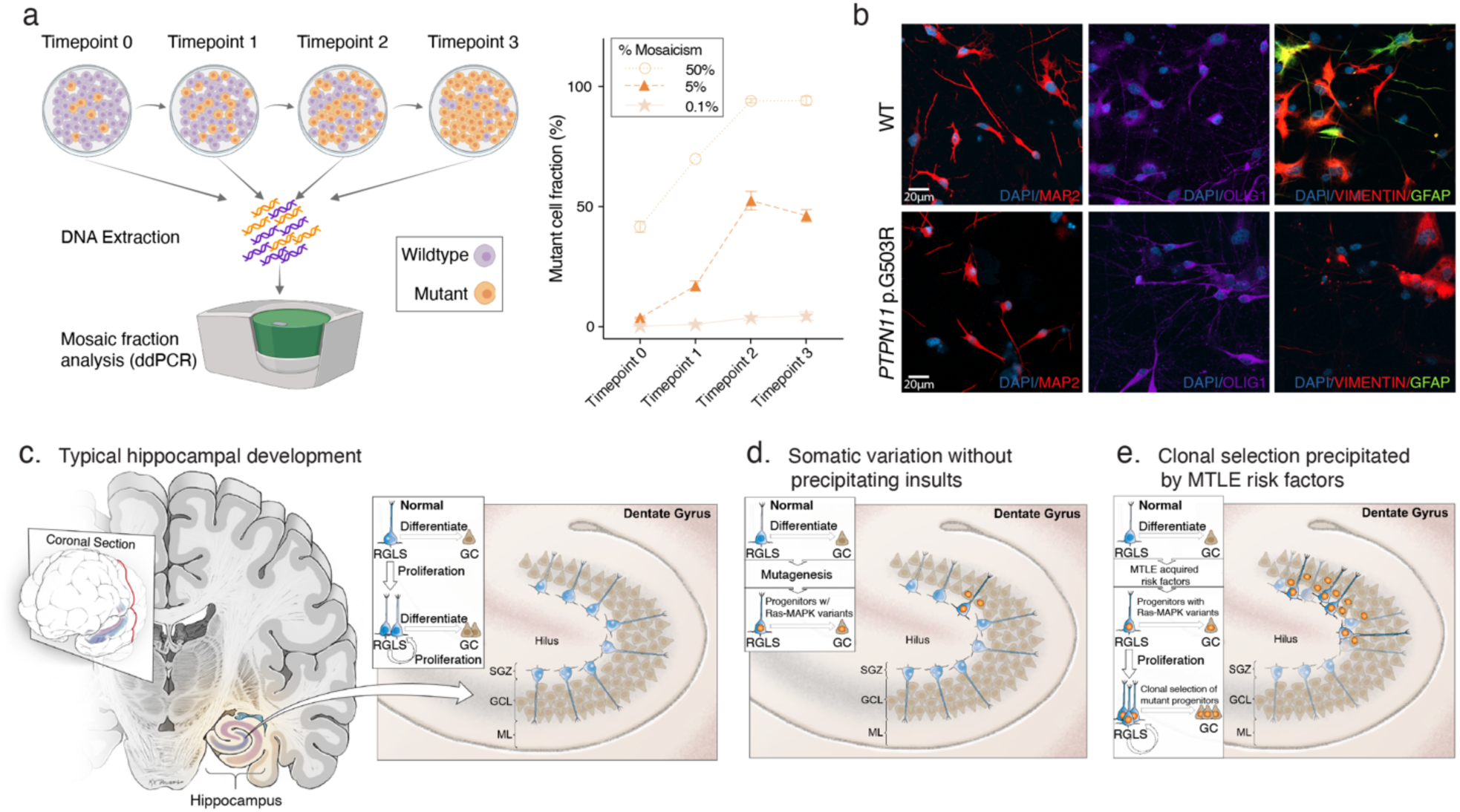
Impact of Ras-MAPK variants on clonal selection and potential contribution in MTLE. (a) Experimental strategy for evaluating clonal competition in a mosaic culture: a wildtype iPSC line mixed with its isogenic engineered PTPN11 p.G503R mutant clone. Changes in the fraction of mutant cells at different timepoints for three mosaic cultures at 1:1,000, 1:20, and 1:1 (MUT:WT) ratios (n = 4 replicates); Mean±SEM is shown. (b) Pseudocolored immunofluorescence images of neurally differentiated WT and MUT iPSCs. DAPI: nuclei, MAP2: neurons, OLIG1: oligodendrocytes, Vimentin and GFAP: astrocytes. (c-e) Proposed model for clonal selection mediating MTLE pathogenesis. During typical development (c), a fraction of neural progenitors are retained in the DG after birth, but neurogenesis declines over time. In the absence of precipitating insults (d), developmentally-acquired somatic Ras-MAPK variants in DG progenitors give rise to inconsequential numbers of mutant neuroglial cells during typical development. However, MTLE risk factors (e) may cause aberrant proliferation and neurogenesis in the DG, which favors the mutant clones with a selective advantage, and may affect hippocampal cellular composition and function long term. RGLSC: radial glia-like stem cell, GC: granule cell, SGZ: subgranular zone, GCL: granule cell layer, ML: molecular layer.

## Discussion

In this multi-center, international, case-control, duplex sequencing study, we identify deleterious somatic Ras-MAPK variants in >40% of MTLE hippocampi with significant enrichment relative to controls, providing strong evidence of the role of these mosaic variants in MTLE pathogenesis. This is the largest cohort of MTLE surgical tissue studied for somatic variants to date and identifies >160 novel, likely deleterious Ras-MAPK somatic variants in 21 genes in 188 MTLE patients, most of which have not been previously described in association with epilepsy. By also sequencing known epilepsy-associated PI3K-mTOR-FCD and DEE genes, we show that enrichment of Ras-MAPK genes is a selective, disease-relevant finding and cannot be simply attributed to increased detection sensitivity from duplex sequencing. The high fraction of variant-carrying cases in a cohort limited to dysplasia- and neoplasia-free hippocampal tissue indicates that somatic Ras-MAPK mosaicism is a common and perhaps defining feature of MTLE with HS, which is supported by the sensitivity analysis predicting a potentially even higher number of somatic Ras-MAPK variants.

Increased variant detection sensitivity and accuracy using duplex sequencing technology facilitated the discovery of many rare somatic variants present in <1% of cells, illustrating that most Ras-MAPK variants have extremely low VAFs, consistent with having arisen relatively late in development. In that regard, Ras-MAPK variants in MTLE differ from PI3K-mTOR variants in FCD type II that likely arise mid-gestation during corticogenesis^24,43,68^, and typically present with higher VAFs^18,20^. Extensive neuronal loss in HS, likely depleting some variant-positive cells, may partly account for the low VAF, but this is unlikely to be the only contributing factor because most variants are also absent in neighboring neocortex. The increasing frequency of somatic Ras-MAPK variants with later age at surgery suggests that somatic variants either arise or undergo clonal proliferation postnatally, and may account for the fact that—unlike FCD—MTLE is most commonly diagnosed in adolescents and adults^69^. Our experimental data demonstrating the strong effect of Ras-MAPK variants on clonal selection, even at low mosaic fractions, are consistent with that observation and with the known role of Ras-MAPK variants in clonal selection^70–73^. Human and mouse data for FCDs suggest that the minimum proportion of mutant cells required for epileptogenesis is extremely low (<0.1%)^74^, supporting the potential functional impact of somatic Ras-MAPK variants in MTLE despite their low VAFs.

Correlation of clinical data with the genetic findings showed strong association between deleterious somatic Ras-MAPK variation and HS, only seen in a handful of previously reported cases^7,8,25^, providing a potential unifying biological mechanism for this common MTLE-associated pathology. On the other hand, the low frequency of PI3K-MTOR-FCD variants in HS contrasts to extratemporal FCD IIb, where the PI3K-MTOR-FCD variants are found in as many as 80% of cases with the most sensitive methodology^75^. The high specificity of the association between lesion type and pathway involved may inform a causal relationship. While sparse and inconsistent documentation of risk factors in clinical notes resulted in no significant associations between MTLE risk factors and hippocampal somatic mosaicism, patients with a history of TBI showed a trend towards increased number of somatic variants. Furthermore, higher deleterious somatic variant count was identified as a negative prognostic indicator of epilepsy surgery. These data suggest that Ras-MAPK somatic variants in MTLE may be associated with specific clinical variables and may inform clinical care, although future prospective studies are needed to confirm these findings.

Somatic Ras-MAPK variants in MTLE are predicted to increase signaling through GOF of activators and LOF of the repressors, which is analogous to other focal epilepsies associated with cancer-associated pathways^16–18,20–23^. The increased enzymatic activity of all tested *PTPN11* variants is consistent with experimental data in human neurons and rodent brains showing Ras-MAPK-activating variants result in cell autonomous neuronal hyperexcitability and spontaneous recurrent seizures, respectively^9,10^. There is also robust clinical evidence from humans that pathologic overactivation of the Ras-MAPK pathway in RASopathies and LEATs can give rise to drug-resistant epilepsy^7,21–23,76^, and that Ras-MAPK inhibition may directly suppress seizures^77,78^. Given the disproportionate loss and increased expression of Ras-MAPK genes in excitatory neurons in our snRNA-seq data, it is likely that Ras-MAPK overactivation in neural cells is a key aspect of MTLE pathogenesis and may also provide a therapeutic rationale for targeting this pathway to treat seizures.

Enrichment of somatic Ras-MAPK variants in the hippocampus relative to the adjacent temporal neocortex, detection of variants in cells from the neural lineage in MTLE in contrast to microglia in AD, and their potential involvement in clonal selection, implicate hippocampal neural progenitors as a likely source of these variants and a potential contributor to MTLE pathogenesis. While neocortical neurogenesis ceases prior to birth^79^, a pool of neural progenitors persists in the DG postnatally that may produce newborn neural cells for at least a few years and perhaps into adulthood^53–62^ (Fig 6c). Our data confirm other reports that somatic variants, including Ras-MAPK variants, are not uncommon in neurotypical brain tissue^80^, suggesting that a few DG progenitors may normally acquire and harbor such somatic variants. In typical development, postnatal DG neurogenesis declines after birth^54,59,60^, so mutant progenitors may have an inconsequential role on hippocampal cellular composition and function (Fig 6d). In the presence of MTLE risk factors, such as early life seizures or head trauma however, experimental models show aberrant proliferation and neurogenesis in the DG^51,64–67^. These early life insults could give the Ras-MAPK mutant progenitors a proliferative and/or survival advantage to undergo disproportionate clonal expansion, starting a cascade of events that ultimately give rise to MTLE (Fig 6e Thus, in some patients the deleterious variants or precipitating insults may be sufficient to cause MTLE, while in others epileptogenesis may require a combination of both genetic and acquired factors. This model may potentially explain how seizures themselves increase the likelihood of future seizures, as is seen with secondary hippocampal involvement in bitemporal epilepsy or in patients with pre-existing neocortical epilepsy^81–83^, given the known effect of seizures to induce DG proliferation^51^. The potential presence of Ras-MAPK mutant clones may also be important in interpreting studies of adult human hippocampal neurogenesis—which occasionally use hippocampal specimens removed for the treatment of epilepsy^53,54^—and in which there is persistent variability in observations between studies performed with similar methods^53–62,84^.

This study provides the strongest human genetic evidence to date on the potentially causal role of somatic Ras-MAPK variants in MTLE with HS and outlines a mechanistic framework for how these variants may be involved in epileptogenesis in the hippocampus. MTLE is however complex and many factors such as the developmental timing of a mutation, the cellular lineage of variant-carrying cells, contribution from germline genetic background, and acquired factors all contribute to disease pathogenesis. Future experimental studies will be needed to specifically test the validity of these assertions and hypotheses in model systems.

## Methods

### Study participants and procedures

Patients with a clinical diagnosis of drug-resistant MTLE at 8 epilepsy surgical centers in North America, Europe, and Australia, who underwent complete or partial anterior temporal lobectomy with hippocampal resection were included in the study. Frozen surgical MTLE surgical tissue was collected and stored at the European Epilepsy Brain Bank (EEBB, Erlangen, Germany), Pitié-Salpêtrière Hospital (PSH, Paris, France), Toronto Western Hospital (TWH, University Health Network, Toronto, Canada), Austin Hospital (AH, Heidelberg, Australia), The Royal Melbourne Hospital (RMH, Parkville, Australia), Yale-New Haven Hospital (YNH, New Haven, USA), Brigham and Women’s Hospital (BWH, Boston, USA), Boston Children’s Hospital (BCH, Boston, USA). Surgical samples and/or extracted DNA were shipped to Boston Children’s Hospital for all downstream processing, sequencing, and analysis. Written informed consent was obtained from all patients or their guardians with local Institutional Review Board or Ethics Committee approval. All research performed with samples obtained from patients was approved by the Institutional Review Board at BCH.

Inclusion criteria included a presurgical diagnosis of MTLE and presence of surgically resected hippocampal tissue with or without hippocampal sclerosis (HS). Patients of all ages, sexes, and demographics were included in the study and no specific selection was made based on these factors. Cases with any history or histopathological evidence of brain tumors including low-grade epilepsy-associated tumors (LEATs) and focal cortical dysplasia (FCD) type IIIB were excluded. Cases with a history of extra-hippocampal abnormalities—such as FCD type I, FCD type II, FCD type IIIA, and periventricular nodular heterotopia—were included only if no dysplastic features were seen in the hippocampus on histopathology. ILAE definitions were used for histopathologic classification of epilepsy-associated lesions^2,85^.

Neurotypical postmortem control tissue was obtained from the NIH NeuroBioBank (the University of Maryland Brain and Tissue Bank, the University of Miami Brain Endowment Bank, the University of Pittsburgh Brain Tissue Donation Program, the Human Brain and Spinal Fluid Resource Center [Sepulveda]) and the European Epilepsy Brain Bank. Control subjects without a documented history of epilepsy and other neurological diagnoses and available frozen hippocampal tissue, were chosen based on age and sex to closely match the MTLE study population. Alzheimer’s disease hippocampal tissue with histopathological evidence of HS was obtained through the NIH NeuroBioBank and Massachusetts Alzheimer’s Disease Resource Center (ADRC).

### Collection and processing of clinical information

De-identified clinical and surgical information was obtained from patients’ charts and summarized by the authorized study team at each site. When possible, the documented radiological and/or histopathologic diagnosis were confirmed again specifically for this study. All clinical information from different sites were organized and codified for consistency and downstream analyses.

### Confirmation of histopathologic and radiological findings

The original clinical histopathological slides for all the cases at EEBB, TWH, BWH, and BCH were reviewed again for this study by expert neuropathologists to confirm absence of tumor and dysplasia. No changes to the original histopathologic diagnoses were made as the result of this analysis. Additionally, since the terminology used to describe hippocampal abnormalities in MTLE is frequently inconsistent and institution-dependent, the ILAE histopathological classification of epilepsy-associated lesions were adopted^2,85^. If the pathology slides were reviewed again, HS was re-classified according to the latest ILAE definition: HS1 (neuronal loss and gliosis in most hippocampal subfields, most severe in CA1 and CA4), HS2 (CA1 predominant neuronal loss), HS3 (CA4 predominant neuronal loss), no-HS (no significant neuronal loss). If HS was present but the hippocampus was only partially viewed or the slides were suboptimal for complete classification, HS-not otherwise specified (HS-NOS) was used. HS-NOS was also used in cases where hippocampal neuronal loss and gliosis were mentioned in the clinical report, but the slides could not be reviewed again for this study.

MRI images were reviewed for all the cases at TWH, BWH, and BCH to evaluate radiological evidence of hippocampal sclerosis. When available, axial and coronal T1- and T2/FLAIR-weighted images of the surgically resected hippocampus on the pre-resection MRI were evaluated for hippocampal atrophy, loss of internal architecture in the hippocampus, and increased hippocampal T2/FLAIR hyperintensity. Presence of any of these MRI findings was interpreted as radiological evidence of hippocampal sclerosis^86^.

### Genomic DNA extraction and QC

Genomic DNA (gDNA) from fresh-frozen surgical and postmortem samples was extracted using EZ1&2 DNA Tissue Kit (Qiagen), Chemagic DNA Tissue 100 H24 kit (Perkin Elmer), or All Prep DNA/RNA kit (Qiagen) according to the manufacturer protocols. All gDNA samples were purified using 2X AMPure XP beads and eluted in low-EDTA TE buffer. Standard quality and purity assessments were conducted using genomic tape (TapeStation, Agilent). All the gDNA samples were quantified using Qubit or PicoGreen (ThermoFisher Scientific) for proper assessment of double stranded DNA concentration.

### Panel design and targeted duplex sequencing

For hybridization capture, probes targeting the exons and exon-intron junctions of 71 genes in epilepsy-associated groups (Extended Data Fig 1c) were designed using Agilent’s SureDesign tool. The list of targeted genes was manually curated to include canonical Ras-MAPK pathway genes from the Broad Institute’s publicly available Molecular Signatures Database (MSigDB), top PI3K-mTOR and FCD-associated genes reported in focal epilepsy^18,20,87^, and top germline developmental and epileptic encephalopathy genes^30^. Genes with the highest published evidence of known pathogenic variation were prioritized. A total of 8129 probes with a genomic size of ∼286.9 kbp were designed and synthesized.

DNA libraries for sequencing were generated by mechanical shearing of 50-200ng of high-quality gDNA followed by library preparation using the Agilent SureSelect XT HS2 DNA Reagent Kit according to manufacturer protocols. The XT HS2 probes included dual indices and duplex MBCs that were used for generating high-quality, error-corrected duplex consensus reads. Duplex MBCs are unique molecular identifiers (UMIs) that are incorporated during adapter ligation to mark individual DNA molecules and facilitate error-correction through consensus between paired DNA strands. The prepared libraries were pooled and sequenced using three Illumina NovaSeq 6000 S4 flow cells with 150 bp paired-end reads.

### Alignment, pre-processing, and variant calling

FASTQ files were trimmed using Trimmer v2.0.3 within AGeNT v2.0.5 (Agilent). The trimmed FASTQ files were aligned to the GRCh38 reference genome using BWA-MEM with the -C option.

Molecular barcode information was added to the aligned BAM files using Locatit within AGeNT v2.0.5 with the following options: -U -S -v2Only -d 1 -m 1 -q 20 -Q 10. Duplicate reads were marked using Picard v1.138, and INDEL realignment was performed using Genome Analysis Toolkit (GATK) v3.6^88^. Duplicate reads sharing the same UMI were collapsed into single-strand consensus reads, with the number of duplicates marked in the XI tag. Two BAM files were generated by requiring at least two duplicate reads (XI≥2 BAM) or one duplicate reads (XI≥1 BAM) for each duplex consensus read.

Somatic single nucleotide variants (SNVs) were called using MosaicHunter v1.0^89^ from both XI≥2 and XI≥1 BAM files as previously described^31^. Potential sSNV candidates were first identified using NaiveCaller with the options -P 0 -q 20 -Q 40. Reads corresponding to these candidate variants were subjected to pileup (-s -Q 0 -q 0) using Samtools, followed by additional filtering with PileupFilter in MosaicHunter using the parameters –minbasequal=20 –minmapqual=40 –asciibase=33 –filtered=1.

Somatic insertion-deletions (INDELs) were identified using Pisces v5.3^90^ and Mutect2^91^. Pisces was run with the parameters: -MinVF 0.0005 –SSFilter false -MinBQ 20 -MaxVQ 100 -MinVQ 0 - VQFilter 20 -CallMNVs False -RMxNFilter 5,9,0.35 -MinDepth 5. Mutect2 was run with default parameters, and only indel variants were extracted. INDEL calls were merged from both Mutect2 and Pisces.

Germline variants were called only on the XI≥2 BAM file using HaplotypeCaller in GATK v3.6 with the options: -l INFO -rf BadCigar -ploidy 2.

### Stringent and sensitive somatic variant call sets

Somatic variants were classified into two categories: “stringent” and “sensitive”. The stringent call set was generated from XI≥2 BAM files of samples with ≥500X depth, followed by variant calling, germline and false-positive filtering, and pathogenicity annotation (XI≥2 calls). This highly accurate call set, produced through uniform and rigorous filtering steps, was used for all the downstream analyses in MTLE, AD, and control samples, except for the analyses shown in Extended Data Fig 4. Importantly, all the case vs control analyses were exclusively performed using the stringent call set.

In contrast, the sensitive call set aimed to maximize the number of deleterious somatic variants. Therefore, in addition to the variants in the stringent call set, XI≥2 calls from low-depth samples (<500X) and experimentally validated XI≥1 MTLE calls (only V and LV variants) were incorporated. The XI≥1 control calls were included in the dataset without orthogonal validation. All the downstream annotation steps were the same for the stringent and sensitive callsets and the details are described below.

For the stringent call set, samples with an average sequencing depth below 500X were excluded from further analysis. 442 out of 462 MTLE samples, 73 out of 74 control samples, and all 17 Alzheimer’s samples met this criterion and were included in the stringent call set. For the expanded call set, all 462 MTLE samples were used for analysis.

### Somatic variant filtration by call sets

Raw somatic variant calls from XI≥2 and XI≥1 BAMs were annotated using ANNOVAR to facilitate the removal of germline and false-positive variants and to assess pathogenicity. Gene-based annotation was performed using refGene^92^, and filter-based annotations included data from gnomAD v4.0 (genome and exome)^93^, dbSNP151_common^94^, COSMIC v70^33^, ClinVar v20221231^39^, and REVEL^95^. Additionally, annotations from the Evolutionary model of Variant Effect (EVE^95^) for missense variants and SpliceAI v1.3.1^96^ for splicing variants were manually appended to evaluate pathogenicity. For EVE, both the EVE score and the pathogenicity classification (by confidence level) were added for each variant. For SpliceAI, the maximum values of DS_AG (delta score of acceptor gain), DS_AL (acceptor loss), DS_DG (donor gain), and DS_DL (donor loss) were reported.

For somatic XI≥2 calls, germline and false-positive variants were filtered using the following criteria: (1) variants with a gnomAD v4.0 genome or exome AF > 0.001, or those overlapping with dbSNP151_common; (2) variants with a VAF > 0.32; (3) variants identified as multi-nucleotide variants (MNVs); (4) variants that appeared in more than two additional samples in XI≥1 calls compared to XI≥2 calls suggesting an error-prone region; (5) variants present in more than 3% of the 532 samples.

For somatic XI≥1 calls, the same filtration criteria were applied as for XI≥2 calls, except that step (4) was omitted and step (5) was modified to exclude variants present in more than 5% of the 553 samples (MTLE, Control, and AD combined). Additionally, manual inspection in Integrative Genomics Viewer (IGV^97^) was performed to validate the variants, which were included in the final XI≥1 calls if validated.

### Germline variant filtration

Raw germline variant calls from XI≥2 and XI≥1 BAMs were annotated as described above. The same set of filters as somatic XI≥2 and XI≥1 calls were applied, except that steps (2) and (4) were omitted and step (5) was modified to exclude variants in more that 3% of the 532 samples.

### Enrichment of deleterious variants

Likely damaging and function altering (i.e., deleterious) variants were identified using a set of criteria based on variant characteristics, public clinical databases (ClinVar and COSMIC), and computational predictions: 1) Variants resulting in stop-gain, frameshift deletions, or frameshift insertions that are likely to disrupt gene function were annotated as deleterious; 2) Variants reported in ClinVar as Pathogenic, Pathogenic/Likely pathogenic, Likely pathogenic, Pathogenic|drug_response|other, or Likely pathogenic|other were considered deleterious; 3) Variants observed more than five times in the COSMIC database showing recurrence were similarly classified as deleterious; 4) Variants were considered deleterious if any of the following criteria were met based on *in silico* predictions on missense and splicing variants from the computational tools EVE, REVEL, and SpliceAI: a) Pathogenic in EVE classification of Class75, Class80, or Class90 per each gene; b) REVEL score greater than 0.6; c) SpliceAI score greater than 0.8. However, if ClinVar classified a variant as benign, likely benign, or benign/likely benign, it was treated as benign, even if computational predictions indicated pathogenicity. This pathogenicity enrichment approach was applied to both somatic and germline variants and is largely based on the filtration strategy we previously developed and benchmarked^7^.

### Mutational spectrum and signature analysis

Mutational spectrum and signature analyses were performed using SigProfiler^98^. All somatic variants were collected, and duplicate entries were removed based on genomic position. Input matrices were generated using SigProfilerMatrixGenerator with targeted regions. Mutational signatures were analyzed using SigProfilerAssignment with COSMIC version 3.4, restricted to exonic regions.

### Variant burden analysis

For all burden analyses for somatic and germline variants, the variant count was normalized by the corresponding sequencing coverage. Coverage was defined as the top 90% of target regions ranked by sequencing depth to exclude regions with excessively low coverage. Variant burden was calculated by dividing the variant count by this coverage for each sample.

To further evaluate the association between variant count and disease type while controlling for additional covariates, a linear mixed-effects regression model was employed. Disease type and covariates including age, sex, and sequencing coverage were modeled as fixed effects, and sequencing batch was included as a random effect. For the age covariate, age at the time of surgery was used for MTLE patients, and age at death was used for controls. Coverage was z-score normalized before inclusion in the model. The linear mixed-effects regression model was formulated as: y_i_ = ɑ_i_ + β_i_ + γ_i_ + δ_i_ + U_i_, where y_i_ is the variant count for sample i, ɑ_i_ represents disease type, β_i_ is age, γ_i_ is sex, δ_i_ is coverage, and U_i_ is batch information as a random effect. This model was fitted using the lme4 package^99^, and statistical significance was assessed using lmerTest.

### Somatic variant selection analysis

Variant selection from somatic evolution was inferred using dNdScv^37^. dNdScv assesses the ratio of non-synonymous to synonymous variants while accounting for the trinucleotide context to identify variants that are under positive clonal selection. The global dN/dS ratio was calculated using the dndscv function with “RefCDS_human_GRCh38_GencodeV18_recommended.rda” as the reference database. Gene-level dN/dS ratios were calculated using geneci, and site-level dN/dS ratios (for hotspot analysis) were calculated using sitednds with the theta option set to maximum likelihood estimates (MLE).

### In-silico simulation of variant detection sensitivity

To simulate somatic variant detection sensitivity, sequencing depth distributions were set at 400×, 600×, 800×, 1,000×, 1,200×, 1,400×, and 1,600×, and VAF distributions were set at 0.1%, 0.2%, 0.4%, 0.6%, 0.8%, 1.0%, 5.0%, 10%, and 20%. A total of 100 sample pairs were randomly selected at SNP loci, where one sample contained a homozygous germline variant and the other sample had the corresponding reference sequence reported in the dbSNP database. Both samples were required to have a minimum read depth exceeding 1600× at each SNP position. Reads located within 1 kb of each SNP were extracted prior to generating mixed synthetic BAM files. Mixed synthetic BAM files, representing each combination of VAF and sequencing depth, were created by mixing two down-sampled BAM files from the original samples. Downsampling ratios were computed using the following formulas, where sample1 represents the sample containing the homozygous germline variant, sample2 is the sample with the reference sequence at the SNP loci, Depthsample denotes the observed read depth at each SNP locus in each sample, and VAFexp and Depthexp represent the predefined expected VAF and sequencing depth categories, respectively: Downsampling ratiosample1 = ((VAFexp × Depthexp) / 100) / Depthsample1; Downsampling ratiosample2 = (((100 − VAFexp) × Depthexp) / 100) / Depthsample2. Variants were then called from the synthetic BAM files using the somatic variant calling method described above. Detection sensitivity was calculated based on the proportion of detected variants across each sequencing depth category.

### Variant visualization

The overall spectrum and gene-level distribution of deleterious variants in MTLE and control samples was visualized using the oncoplot and lollipop functions in R maftools v2.22.0^100^.

### Correlation of variants and clinical variables

Mean deleterious variant counts were computed for each clinical variable in the MTLE cohort. To evaluate statistical significance, an equal number of samples per clinical variable were randomly selected, and the mean deleterious variant count was recalculated across 1,000 iterations. P-values and 95% confidence intervals were derived from the resulting empirical distribution.

Associations between variants in each pathway and clinical variables were assessed using Fisher’s exact test on 2×2 contingency tables, defined by the presence or absence of deleterious variants and the corresponding clinical category. Early and late onset ages were defined as <5 years and >20 years, respectively, while early and late surgery ages were defined as <12 and >35 years, respectively. The e-e, e-l, and l-l groups included 62, 30, and 64 samples, respectively.

Decile groups for onset, surgery, and death ages were generated from MTLE samples with available clinical data, resulting in 39, 43, and 7 samples per decile group, respectively.

### Orthogonal validation of candidate somatic variants

Candidate somatic variants from the stringent and sensitive call sets were experimentally tested for validation using amplicon sequencing and/or ddPCR based on our published protocol^7^. Based on the orthogonal validation experiments, the variants were divided into three categories: 1) Validated (V)—if the variant was confirmed with >=2 independent, deeply sequenced amplicons or with ddPCR. 2) Likely Validated (LV)—if the variant was confirmed with only 1 deeply sequenced amplicon. 3) Not Validated (NV)—if the variant was not confirmed with amplicon sequencing or ddPCR.

For amplicon validation, custom primers were designed for each candidate variant using the default settings in Primer3^101^ to generate 150-500bp amplicons. The primers were commercially synthesized (IDT) and tested on human genomic DNA (Promega) to confirm generation of only one amplicon product at the expected size. The validated primers were then applied to 5-20ng of gDNA extracted from brain tissue to generate amplicons for sequencing using the Phusion High-Fidelity PCR Master Mix (Thermo Fisher). The PCR products were cleaned up using 2X AMPure XP beads (Beckman Coulter), and run on a gel to confirm the presence of the desired amplicon product. Different amplicons were pooled together and Illumina sequenced with target minimum sequencing depth of 10,000 reads per each unique amplicon. The raw reads were aligned to the reference genome (hg38) using BWA v0.7.17 and then visualized on IGV to confirm the presence of each candidate variant and to ascertain that the variant is not in an error-prone region. VAF was calculated based on the total number of reference (REF) and alternate (ALT) alleles. VAFs < 0.1% were considered as NV since they could not be reliably distinguished from sequencing and experimental artifacts.

For ddPCR validation, commercially available and validated TaqMan probes (Thermo Fisher) with FAM and VIC fluorescent markers were used for detection of specific REF and ALT alleles. First, 5-10ng gDNA was digested using the HindIII restriction enzyme. Then a reaction mix was prepared for each variant using the digested gDNA, 2X ddPCR Supermix for Probes (no dUTP) (Bio-Rad), and the variant-specific specific TaqMan probe set. All reactions were performed in two technical replicates. The reaction mixture was incubated at 37°C for 10 minutes and loaded into cartridges for partitioning and emulsification of the reaction in droplet generation oil for probes (Bio-Rad) in a QX200 Droplet Generator. Following droplet generation, the emulsified droplets were transferred to a 96-well plate for signal amplification in a thermal cycler using the following program: 95°C for 10 mins (1 cycle), 94°C for 30 sec followed by 60°C for 1 min (40 cycles), 98°C for 10 mins (1 cycle), cool down 4°C. After signal amplification, the number of FAM/VIC positive and negative droplets was quantified on a QX200 droplet reader (Bio-Rad). The ddPCR raw data was analyzed using QuantSoft v1.0596 (Bio-Rad), and the VAFs were calculated based on the total number of REF and ALT alleles.

### Single-nucleus RNA sequencing (snRNA-seq) experimental protocol

Nuclei isolation and droplet-based snRNA-seq are based on previously published protocols^43,102^ with minor modifications that are described below. Approximately 5-20 mg of frozen hippocampal tissue from 6 surgically resected MTLE cases and 2 neurotypical controls from the NIH NeuroBioBank were finely minced using a chilled sterile scalpel at -20C and transferred to a clean Potter-Elvehjem Tissue grinder containing 3mL of homogenization buffer (0.25M Sucrose, 5nM MgCl2, 25mM KCl, 10mM Tris-HCl, 1µM DTT, 0.2U/mL RNase Inhibitor), and homogenized thoroughly. The resulting homogenate was filtered through a 40µm cell strainer and centrifuged at 500 rcf for 5 mins to form a nuclear pellet. The nuclear pellet was resuspended in 500uL of immunostaining buffer (5nM MgCl2, 1% BSA, 0.2U/mL RNase Inhibitor, in 1X PBS) and then stained with DAPI (1:500; Thermo Fisher Scientific, D1306) for 10 mins prior to proceeding to fluorescence activated nuclei sorting (FANS).

Nuclei sorting was performed on a FACS Aria II cell sorter equipped with BD FACSDiva software at the BCH HSCI flow cytometry core, selecting all DAPI-positive nuclei (SSC-A and FSC-A gate boundaries were set to minimize doublets). Single-nuclei snRNA-seq was performed using the 10X Genomics Chromium Next GEM Single Cell 3ʹ Reagent Kit v3.1. 13,500 nuclei/reaction were used to load the 10X Chromium, and were processed the same day for gel-bead in emulsion (GEM) generation, barcoding, and cDNA amplification, following manufacturer instructions. Libraries were sequenced (paired-end single indexing) on an Illumina NovaSeq6000 targeting ∼50,000 read-pairs per single nucleus.

### snRNA-seq data preprocessing

We analyzed snRNA-seq data from two sources (all samples are shown in Extended Data Fig 7): 1] data generated for this manuscript (n=8), 2] previously published data from Ayhan et al.^46^ (n=3), representing 5 MTLE cases with HS, 4 MTLE cases without HS, and 3 controls. Gene counts matrices for our data were generating by aligning the raw FASTQ files to the GRCh38 reference human genome using Cell Ranger v7.1.0 (10X Genomics) and the reference file “gex-GRCh38-2020-A” with default settings, including intronic reads. Counts data were obtained from Ahyan et al. 2021 using the Gene Expression Omnibus accession number 160189. Posterior and Anterior MTLE samples from female individuals were used for our analysis, as all our MTLE samples are also from female individuals. For all analyses using both datasets, only genes with counts in both datasets were used.

Seurat v5.3.0^103^ R package was used for all subsequent downstream analyses. Standard QC was performed on both datasets together: 1] percent mitochondrial gene expression <5%, 2] >200 but <10,000 unique genes (*nFeature_RNA*) to exclude doublets and low quality nuclei. High quality nuclei data were then normalized (*NormalizeData)* and scaled (*ScaleData)* according to default Seurat processing parameters. CCA Integration was then performed based on donor identity to avoid donor-specific or dataset-specific clustering. Dimensionality reduction was performed using PCA (*RunPCA)*, and the top 20 principal components were used for downstream analysis based on visual inspection of an elbow plot of the proportion of variance captured by each PC. Upon clustering, more doublets were removed based on expression of marker genes from multiple cell types in the same nuclei. After iterative clustering and doublet removal, the final clustering parameters used were 20 principal components and 0.8 resolution, and all other parameters left as defaults. UMAP (*RunUMAP)* was used for dimensionality reduction and visualization. Annotation was performed manually based on known cluster marker used in our prior publications^43,104^ and the BRAIN Initiative Cell Census Network Human Brain Cell Atlas v1.0^105^.

### Differential cell type composition analysis

To determine whether there were differences in cell type composition between MTLE and controls, first the overall differences in cell type composition were assessed using a chi-squared proportion test (base R *chisq.test*). Then to determine enrichment or depletion of specific cell types in MTLE relative to controls, pairwise cell type proportion statistics was performed using the two-sided Wilcoxon test. The fold difference statistics were done using a permutation test with 1,000 permutations in the scProportionTest R package v0.0.0.9000^106^.

### Gene module expression analysis

To evaluate the expression of genes from the duplex sequencing panel in specific cell types, three gene modules consisting of Ras-MAPK, PI3K-mTOR-FCD, and DEE genes were created in Seurat. The native Seurat *AddModuleScore* function was used with default settings and number of control genes set to 100. Statistical analysis comparing expression of modules across groups (MTLE vs control) was done using Wilcoxon tests. Module expression was considered significantly different between groups if the p value was less than 0.05.

### Differential gene expression and gene ontology analysis

Since snRNA-seq data from Ayhan et al. was sequenced at much lower coverage than the data generated for this paper, Ayhan et al. data was excluded from differential gene expression and gene ontology analysis.

EdgeR v4.4.2^107^ with default parameters was used on sample-bulked counts data for each cell type for all differential gene expression analysis. Benjamini-Hochberg correction was applied to generate FDR values on all nominal p values for genes, and genes were considered significantly changed if the FDR was less than 0.05.

For Gene Ontology (GO) term analysis, all GO terms from the Gene Ontology: Biological Process (GO:BP) and Kyoto Encyclopedia of Genes and Genomes (KEGG) databases^108,109^ were analyzed for enrichment in significant genes in each listed comparison. The R Package gprofiler2 v0.2.3^110^ with default parameters was used to calculate GO term enrichment and significance in differentially expressed gene sets using a cumulative hypergeometric test. Terms with Benjamini-Hochberg-adjusted p-values <0.05 were considered statistically significance. The top 20 terms by significance—or fewer if <20 statistically significant terms available—were plotted in bar plots for each group.

### Statistical analyses of genomic and transcriptomic data

All the statistical analyses of the genomic and transcriptomic data were performed in R, except for the Fisher’s exact test used in Fig 4b which is performed in Python. Two-sided Wilcoxon rank-sum test was used to compare the distributions of non-parametric data (confidence intervals were obtained via random sampling with 1,000 replacements) and Fisher’s exact test was used for comparison of associations between categorial variables. To evaluate the effect of mutational burden, we employed a linear mixed-effects model using the lme4 v1.1 package^99^, and statistical significance was assessed with lmerTest. The 95% confidence intervals and p-values were calculated from the lmerTest package. dN/dS ratio was calculated using the dNdScv R package^111^ and 95% confidence intervals were obtained from the package output; dN/dS ratios with a lower bound greater than 1 were considered statistically significant. For Fig 4a, p-values were derived from the normal distribution of 1,000 bootstrapped means, based on the central limit theorem. For the cumulative sum distribution in Fig 3g, we used the Kolmogorov–Smirnov (K–S) test to calculate p-values. P-values <0.05 were considered statistically significant. In cases where correction for multiple hypotheses testing was performed, adjusted p-value < 0.05 was considered statistically significant.

### Expression and affinity purification of SHP2 proteins

Plasmids encoding wild-type SHP2 residues 1-525 (SHP2^WT^ and SHP2^MUT^) and 200-525 (PTP domain only) with an N-terminal 10x histidine tag separated by a tobacco etch virus (TEV) protease cleavage site were obtained from the lab of Stephen Blacklow^63,112^. Mutant plasmids (expressing SHP2^MUT^: D61N, A72V, E76K, E139D, N308D, G503R, G503V) were synthesized by GenScript (Piscataway, NJ) using site-direct mutagenesis and verified with whole plasmid sequencing. The SHP2 expression and purification protocol was based on prior published work from the Blacklow lab^63^ and is summarized here as well. Transformed BL21 (DE3) E. coli (New England BioLabs) were grown at 37°C in 500 ml Luria Broth containing 100 ug/ml ampicillin until reaching an OD600 of 0.6-0.8. SHP2 expression was induced with 0.5 mM IPTG, and cells were grown overnight at 18°C. Cell pellets were resuspended in 50 ml lysis buffer containing 50 mM HEPES pH 8.3, 25 mM imidazole, 200 mM NaCl, 0.1% Triton X-100, 5% glycerol, 2 mM mM tris(2-carboxyethyl)phosphine (TCEP), and cOmplete EDTA-free protease inhibitor (Roche) and lysed via sonication. Cell lysates were clarified by centrifugation at 20,000g for 30 min, and the supernatant was incubated with 3 ml of HisPur Ni NTA Resin (Thermo Scientific) for 1 hour at 4°C. The resin was washed 3 times with 20 ml wash buffer (lysis buffer lacking Triton X-100) then 3 times with wash buffer containing 50 mM imidazole before transfer to an Econo-Pac chromatography column (Bio-Rad) for elution. SHP2 elution was achieved by the addition of 2 ml wash buffer containing 300 mM imidazole and repeated four times. The most concentrated elution fraction was incubated with TEV protease (1 mg TEV:20 mg SHP2) overnight at 4 °C to remove the tag then diluted 1:1 with wash buffer lacking NaCl followed by injection onto a MonoQ 10/100 column. The column was equilibrated with 50 mM HEPES pH 8.5, 50 mM NaCl, 5% glycerol, and 2 mM TCEP, and protein was eluted with a 10 column-volume gradient from 50 to 500 mM NaCl. Peak fractions containing pure SHP2, as assessed by SDS polyacrylamide gel electrophoresis and Coomassie staining, were pooled and loaded onto a PD-10 desalting column (Cytiva) to exchange the protein into storage buffer (50 mM HEPES pH 7.5, 150 mM NaCl, 5% glycerol, and 2 mM TCEP). Aliquots were snap frozen in liquid nitrogen and stored at -80°C.

### Quantification of basal SHP2 phosphatase activity

To measure basal phosphatase activity, a 1:2 dilution series was prepared for SHP2^WT^, each SHP2^MUT^(D61N, A72V, E76K, E139D, N308D, G503R, G503V), and PTP from 100 nM to 6 pM in the presence of 200 uM DiFMUP in assay buffer (60 mM HEPES pH 7.2, 75 mM KCl, 75 mM NaCl, 1 mM EDTA, 0.05% Tween 20, 2 mM TCEP), and changes in fluorescence emission resulting from DiFMUP (6,8-Difluoro-4-Methylumbelliferyl Phosphate) dephosphorylation were measured at 450 nM with 358 nm excitation on a Spectramax M5 instrument. An enzyme concentration was selected at the center of each linear response over 10 min and used in substrate titration experiments (2.5 nM wildtype; 0.0125 nM PTP, E76K, A72V, and G503R; 0.125 nM D61N and N308D; 0.15 nM E139D; 0.02 nM G503V). To determine basal SHP2 enzyme kinetics, respective concentrations of SHP2^WT^, each SHP2^MUT^, and PTP were prepared in assay buffer and added to a linear titration of DiFMUP, from 1 mM to 500nM, and DiFMUP dephosphorylation was measured on a Spectramax M5 plate reader at 358 nm excitation and 450 nm emission. Raw velocity data were converted to product (DiFMU)/time by preparing a standard curve of 6,8-difluoro-7-hydroxy-4-methylcoumarin (DiFMU)/ DiFMUP and normalized to enzyme concentration to yield relative velocity, which was fit to the Michaelis−Menten equation to extrapolate kinetic parameters: V= Vmax X [S] / (Km + [S]). V represents enzyme velocity, Vmax is the maximal enzyme activity, Km is the Michaelis−Menten constant, and [S] is the substrate concentration. In vitro SHP2 phosphatase assays were performed in replicates of three on the same day and were repeated for a total of three times on different days. The results of a representative experiment are shown in Fig 7D and data from all three independent experiments was included in Fig 7E. All the analyses were performed using GraphPad Prism v10. Two-tailed t-test was used for statistical analysis of catalytic efficiency given expected normal distribution.

### *PTPN11* (SHP2) variant analysis and 3D visualization

The lollipop visualization of *PTPN11* variants over the gene body was generated using ProteinPaint, a web-based visualization tool developed by St. Jude Children’s Research Hospital (https://proteinpaint.stjude.org/).

PyMOL Molecular Graphics System v3.1.5.1 (Schrödinger, LLC) was used to generate 3D structural visualization of SHP2 protein and the variants. Crystal structure of SHP2 was downloaded from the RCSB Protein Data Bank (PDB ID: 2SHP), then rendered in a cartoon representation and colored by domain (N-SH2 domain – amino acids 1-111, C-SH2 domain – amino acids 112-216, and PTP domain – amino acids 217-525). The PTPN11 variants of interest were highlighted in red spheres, without showing the main and side chains. The structural rendering was oriented to provide an optimal view of the variants.

### Cell type-specific variant enrichment analysis

Nuclei isolation was performed as described above under snRNA-seq experimental protocol. The nuclear pellet was resuspended in 500uL of immunostaining buffer (5nM MgCl2, 1% BSA, 0.2U/mL RNase Inhibitor, in 1X PBS) and evenly split into two tubes. An additional 250uL of immunostaining buffer was added to each aliquot. One aliquot was stained with DAPI (1:500; Thermo Fisher Scientific, D1306), SOX10-PE (1:100; Novus Biologicals, NBP2-59621R), NeuN-Alexa-488 (1:500; Millipore Sigma, MAB377X), and AQP4- Alexa-647 conjugated (1:125; Cell Signaling, 89851S) antibodies to identify oligodendrocytes, neurons, and astrocytes, respectively. The second aliquot was stained with DAPI (1:500; Thermo Fisher Scientific, D1306), and CSF1R-PE (1:100; Cell Signaling, 65396S), with 0.1% Triton X for labeling microglia labeling. Nuclei were incubated on a rotator for 60 minutes at 4°C, then washed twice with staining buffer prior to fluorescence activated nuclei sorting (FANS).

Nuclei sorting was performed on a FACS Aria II cell sorter equipped with BD FACSDiva v8.0 at the BCH HSCI flow cytometry core. Cell type specific FANS from human brain tissue was previously published and validated by our lab for neurons^113^, oligodendrocytes^35^, and microglia^31^, and the protocol for sorting astrocytes was specifically developed and validated for this study. FlowJo v10.0 was used to visualize the sorting gates. After gating for DAPI to isolate nuclei, the following FANS gating strategy was used to enrich for specific cell types:

a. Neurons: DAPI+, NeuN+
b. Astrocytes: DAPI+, NeuN-, SOX10-, AQP4+
c. Oligodendrocytes: DAPI+, NeuN-, SOX10+
d. Microglia: DAPI+, CSF1R+

The sorted nuclei were pelleted by centrifuging at 10,000 rcf for 10 mins and the supernatant was removed. gDNA from each sorted nuclei population was extracted using EZ1&2 DNA Tissue Kit Qiagen, 953034). All gDNA samples were purified using 2X AMPure XP beads and eluted in DNase/RNase-free water.

To determine the mutant cell fraction in each sorted population, we performed ddPCR according to the protocol described above, using commercially available TaqMan probes (Thermo Fisher) for known variants for each sample. Only reactions that had >200 droplets with FAM/VIC signal were included in the analysis to minimize bias from under-sampling. To determine enrichment of a specific cell type relative to others, first a pseudobulk VAF was calculated for each sample by dividing the sum of all the ATL+ droplets by all the REF+ droplets for all the cell types. For each sample, sorted cell types with VAF greater that the pseudobulk VAF (i.e., ratio >1) were considered enriched.

### iPSC clonal competition experiment

A wildtype (WT) iPSC line and its engineered isogenic mutant (MUT) harboring the *PTPN11* (SHP2) p.G503R were obtained from the laboratory of Dr. Maria Kontaridis at Masonic Medical Research Institute (Utica, NY). These lines were initially derived from a healthy individual with appropriate informed consents in place and have been previously characterized and published^114,115^. The Institutional Review Board (IRB) at BCH reviewed and approved the use of these iPSC lines.

After initial QC measures including karyotyping to confirm absence of large chromosomal abnormalities and amplicon sequencing to verify the genotype, the WT and MUT iPSC lines were first expanded. For expansion, the cells were thawed into the mTeSR™ Plus media (STEMCELL Technologies) supplemented with 2 µM Thiazovivin ROCK inhibitor (Tocris Biosciences) and 1,000 IU Penicillin/1,000 µg/mL Streptomycin, counted, and plated on Geltrex (Gibco)-coated 15 cm plates. Once cells reached approximately 70% confluency (when colonies started merging), they were passaged and counted for the mixing experiment.

For the mixing experiment, cells were plated in a 24-well TC-treated, Geltrex-coated plate with three experimental conditions, each condition having four replicates: 50% MUT + 50% WT, 5% MUT + 95% WT, 0.1% MUT + 99.9% WT. Specifically, after lifting the cells with Accutase (Sigma-Aldrich), they were resuspended in 1 mL mTeSR™ Plus containing 2 µM Thiazovivin ROCK inhibitor and antibiotics, then counted. From these counts, appropriate volumes corresponding to 50,000 cells per replicate were transferred into separate 50 mL conical tubes to maintain independent replicates. From each replicate’s suspension, 10,000 cells were set aside and cryopreserved in 1 mL mFreSR™ medium at -150°C. The remaining 40,000 cells per replicate were plated into each well with fresh mTeSR™ Plus medium containing Thiazovivin ROCK inhibitor and antibiotics. The Thiazovivin ROCK inhibitor was removed after 24 hours. Upon reaching approximately 70% confluency, cells underwent passaging, counting, and replating again at 40,000 cells per well. This passaging and replating step was repeated three times in total, with 10,000 cells banked at each passage.

To quantify the number of MUT and WT cells at each timepoint (passage), the reference (REF) and alternate (ALT) alleles were quantified using ddPCR. First, the 10,000 cryopreserved cells from the above mixing experiments were thawed and gDNA was extracted using the EZ1&2 DNA Tissue Kit (Qiagen) and quantified with Qubit. Then, the REF and ALT alleles at each timepoint were quantified using ddPCR with the Taqman probe C_322313626_10 (ThermoFisher) according to the protocol described above. Assuming the mutant iPSCs have one REF and one ALT alleles and that the WT cells have two REF alleles, MUT:WT cell fractions at each timepoint were estimated from the ddPCR allele counts using the following formula: MUT:WT cell fraction = ALT/(ALT+(REF-ALT)/2). Timepoint 0 represents the first mix (pre-plating) and timepoints 1-3 represent cells at the time of passages 1-3.

### Neural differentiation of iPSCs and immunocytochemistry

The WT and MUT (*PTPN11* [SHP2] p.G503R) iPSC lines described above were expanded on Geltrex-coated 15 cm tissue-culture plates in mTeSR™ Plus media (STEMCELL Technologies) supplemented with 2 µM Thiazovivin ROCK inhibitor (Tocris Biosciences) and 1,000 IU/mL penicillin-1,000 µg/mL streptomycin. Neural induction was carried by plating 2.0 x 10⁶ iPSCs per well in 6-well, tissue-culture treated plates pre-coated with Geltrex (1:200 in 50 mL cold DMEM/F-12) in STEMdiff Neural Induction Medium with SMADi Supplement, 2 µM Thiazovivin, and antibiotics. The medium was changed the following day to remove ROCK inhibitor, and daily thereafter. NPCs were passaged every seven days with Accutase and banked after the third passage.

For neural differentiation, NPCs were transitioned to the STEMdiff Forebrain Neuron Differentiation Medium (STEMCELL Technologies) and maintained for seven days on Poly-L-ornithine (15 µg/mL) and Laminin (5 µg/mL)-coated plates. Medium was changed daily. At this stage, neuronal precursors were formed and subsequently passaged into BrainPhys Neuronal Medium supplemented with the STEMdiff Forebrain Neuron Maturation Supplement (STEMCELL Technologies). Cells were plated onto Poly-L-ornithine-Laminin-coated 8-well chamber slides (Thermo Fisher) at 5 x 10⁴ cells per well and matured for an additional 23 days with full medium changes every two days.

To identify differentiated cell types using immunocytochemistry, cells were equilibrated to room temperature, washed once in PBS, and fixed with 4% paraformaldehyde for 15 minutes. Slides were washed five times in PBS and blocked for one hour at room temperature in Serum Blocking Buffer (5% normal donkey serum in PBS containing 0.05% Tween-20). Primary antibodies were diluted in blocking buffer and applied overnight at 4°C: rabbit anti-MAP2 (1:200, Cell Signaling #8707), goat anti-OLIG1 (1:500, R&D Systems AF2417), rabbit anti-Vimentin (1:200, Cell Signaling #5741), and mouse anti-GFAP (1:800, Cell signaling #3670). The following day, slides were washed five to six times in PBS and incubated with donkey anti-rabbit IgG (H+L) Alexa Fluor 555 (1:500, Thermo Fisher A31572), donkey anti-goat IgG (H+L) Alexa Fluor 594 (1:500, Thermo Fisher #A11058), and donkey anti-mouse IgG (H+L) Alexa Fluor 488 (1:500, Thermo Fisher A21202) for two hours at room temperature. DAPI (1:1,000, Cell Signaling #4083) was included during secondary incubation for nuclear counterstaining. Slides were washed five times in PBS, air-dried, and mounted with ProLong Diamond Antifade Mountant (Thermo Fisher, #P36961) before storage at 4°C. Fluorescent images were acquired at 40x magnification using a Zeiss LSM880 confocal microscope with Fast Airyscan and processed in FIJI (ImageJ2 v2.14.0) using the same settings for all samples.

### Generating schematics

Schematics in Fig 1a, Fig 2f, Fig 4a, Fig 5b, Fig 5e, and Extended Data Fig 9d were created by BioRender.com and modified in Adobe Illustrator.

Author contributions

SK and CAW conceptualized the study and SK, CAW, AYH, EAL, and ZZ designed the genomic experiments. SK, RBR, FB, VN, BM, TAV, HMC, SG, MSH, KB, JC, APM, JAK, MT, PK, TJO, SFB, IES, PP, EL, JDR, GRC, RAS, AP, RMR, EZE, ND, DDS, CGB, KTK, CJB, and IB recruited patient participants and contributed to MTLE sample collection, processing, and storage. SK, RBR, YC, BC, and IB contributed to neurotypical control sample collection, processing, and storage. KSMB and MM contributed to AD sample collection, processing, and storage. SK, RBR, LM, FB, TAV, SG, MSH, KB, JC, APM, JAK, MT, PK, TJO, SFB, IES, PP, AMD, ND, DDS, KTK, and IB assisted and/or supervised collection of de-identified clinical data from MTLE patients. RC, FB, AFG, SA, IB performed histopathological review and classification of the epilepsy surgical resections. SK, TK, and EY performed radiological review and classification of the presurgical brain MRI. SK performed the targeted duplex panel sequencing experiments and RBR and YC conducted amplicon sequencing and ddPCR for orthogonal validation of candidate somatic variants. SK, MB, YW, and AYH analyzed the panel sequencing data and SMZ and AL manually annotated the germline variants. SK, MB, and LM processed and analyzed the clinical data. SK designed and performed the snRNA-seq experiments. SK and BF analyzed the snRNA-seq data. SK, EDE, AT, and SCB designed the SHP2 proteomics experiments and EDE and AT performed these experiments. SK, EDE, and SCB analyzed and interpreted the proteomics experiments. DP performed gene-level analysis of SHP2 variants and created the 3D model. SK and RR performed the cell type-specific sorting experiments with guidance from ZZ. SK analyzed the cell type-specific sorting experiment results. AER, LNS, and MIK generated and contributed the iPSC lines. AT and RR performed the iPSC clonal competition experiments and AT and SK analyzed the results. AT performed the iPSC neural differentiation and microscopy experiments. SK, MB, RBR, BF, EG, and DP generated all the figures. SK and MB synthesized and interpreted all the data and wrote the paper. CAW, EAL, and SK provided funding for the experiments and the bioinformatic analysis. All the authors edited the manuscript. CAW, EAL, AYH, and SK directed the research.

## Other contributions

We thank the patients and their families for their invaluable donations to the advancement of science. We thank the following individuals for their contributions to obtaining brain tissue for research and assistance with genomic data sharing: Dustine Reich (Brigham and Women’s Hospital), Julie Lerond (La Pitié-Salpêtrière Hospital), Mandana Movahed (University Health Network), Rebeca Borges Monroy (Boston Children’s Hospital), Javier Ganz, PhD (Boston Children’s Hospital), Jennifer E. Neil, MS, CGC (Boston Children’s Hospital), and Robert Sean Hill (Boston Children’s Hospital). We thank Kenneth Probst (Xavier Studio) for the illustration in Extended Data Figure 10. We thank Ronald Mathieu, PhD at BCH-HSCI flow cytometry Facility and Rajesh Krishnan at BWH ARCND flow cytometry core for their assistance with fluorescence assisted nuclear sorting. We thank IDDRC Molecular Genetics Core Facility at Boston Children’s Hospital (supported by NIH award P50HD105351) for providing instruments and support for ddPCR and snRNA-seq experiments. We thank Boston Children’s Hospital Core Repository for Neurological Disorders, European Epilepsy Brain Bank, NIH NeuroBioBank (the University of Maryland Brain and Tissue Bank, the University of Miami Brain Endowment Bank, the University of Pittsburgh Brain Tissue Donation Program, the Human Brain and Spinal Fluid Resource Center [Sepulveda]), and Massachusetts General Hospital MIND brain bank for providing tissue for this study.

## Data availability

All the panel sequencing and snRNA-seq data generated for this study is currently being deposited in the database of Genotypes and Phenotypes (dbGaP) and will soon be available through the following accession number: phs004124.v2.p1.

---

**Extended Data Figure 1.**
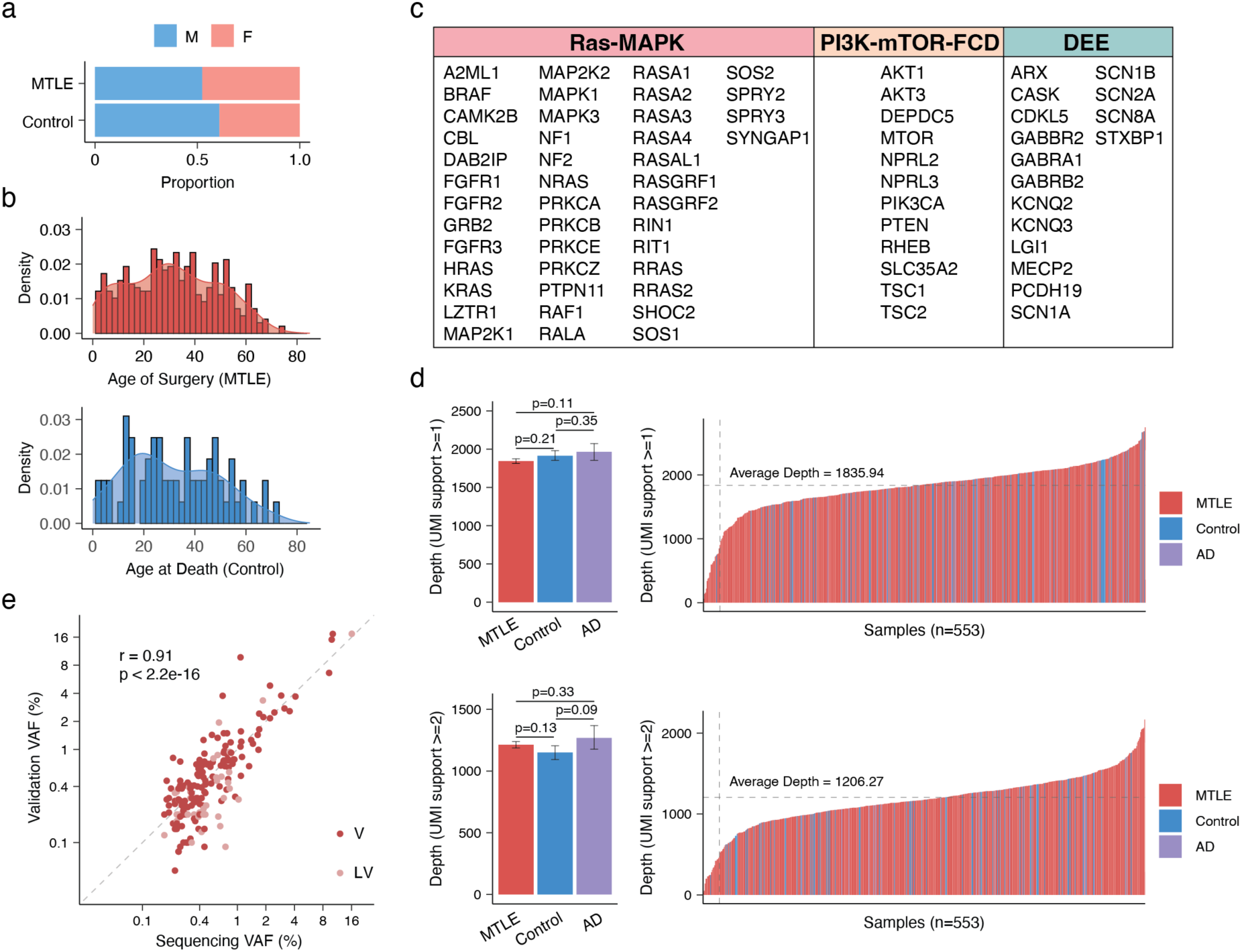
Sequencing parameters and sample demographics. (a) Sex distribution in MTLE and neurotypical control groups. (b) Age distribution at sample collection—age at the time of surgery for MTLE and age at death for neurotypical controls. (c) List of genes in the custom panel. (d) Mean sequencing depth distribution across groups based on UMI support thresholds (Two-sided Wilcoxon test; 95% CI via 1,000 bootstrap iterations). Right panels show per-sample sequencing depth distribution. (e) Correlation between VAFs from targeted sequencing and orthogonal validation experiments (n = 205). The correlation coefficient (r) and p-value were calculated using the Pearson correlation test.

**Extended Data Figure 2.**
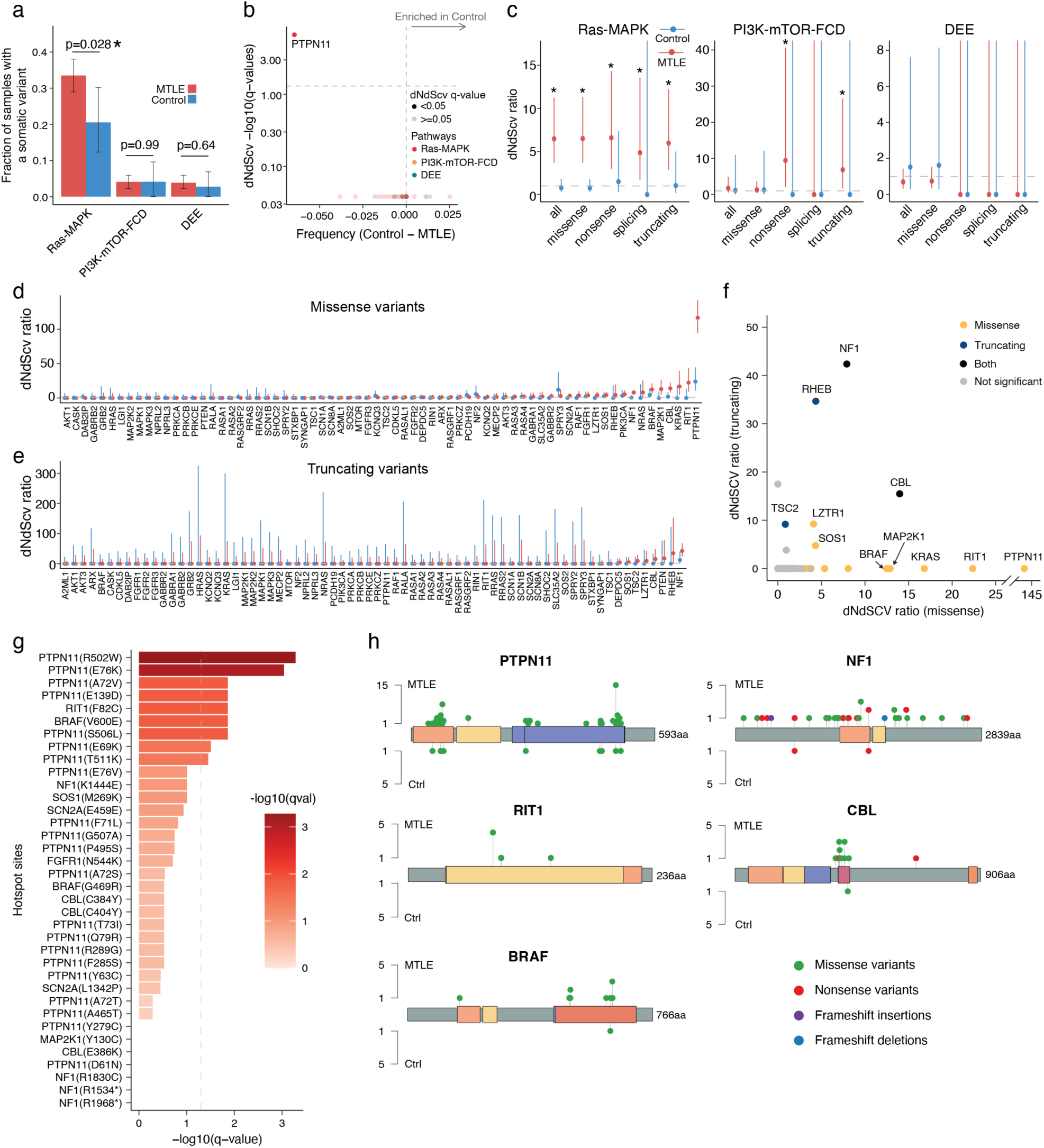
Positive selection of Ras-MAPK gene variants in MTLE. (a) Proportion of samples with deleterious somatic variants in each gene group (Two-sided Wilcoxon test; 95% CI via 1,000 bootstrap iterations). (b) Gene-level dN/dS analysis for control variants. Genes to the right of the vertical dashed gray line are enriched in control versus MTLE and genes above the horizontal dashed gray line show significantly increased nonsynonymous to synonymous variant ratios in controls (q-value < 0.05). (c) dN/dS ratios of Ras–MAPK, PI3K-mTOR-FCD, and DEE-related genes, categorized by variant type. (d-e) Gene-level dN/dS ratios for missense (c) and truncating (d) variants in MTLE and control groups. Lower bound of the 95% CI > 1 indicates positive selection. (f) Summary of gene-level dN/dS ratios for MTLE missense and truncating variants. Positively selected genes are color-coded by variant type. (g) –log10(q-value) of site-level dN/dS scores, highlighting MTLE mutational hotspots ranked by significance. Dashed gray line demarcates q-value = 0.05. (h) Lollipop representation of deleterious somatic variant distribution across the gene body. Only the most strongly selected genes (*PTPN11*, *NF1*, *RIT1*, *CBL*, and *BRAF*) are shown.

**Extended Data Figure 3.**
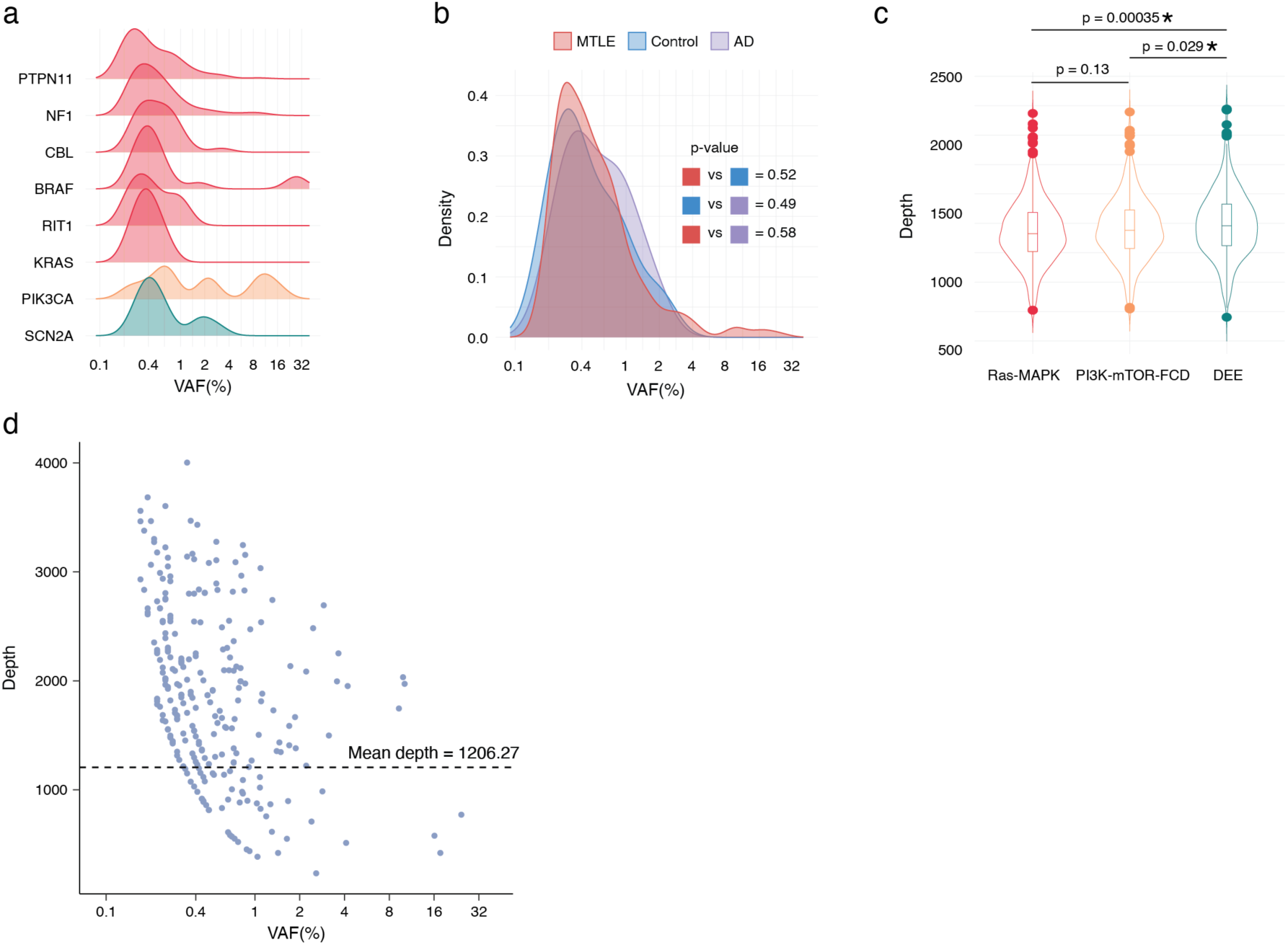
VAF distribution across cohorts and gene groups. (a) VAF distribution of representative genes from the Ras-MAPK, PI3K-mTOR-FCD, and DEE gene groups. (b) Distribution of deleterious somatic variant VAFs by sample cohort (Two-tailed Wilcoxon test; 95% CI via 1,000 bootstrap iterations). (c) Sequencing depth per gene group (UMI>=2; two-sided Wilcoxon test). Boxes show median±25% quartiles; asterisks indicate statistical significance. (d) Association between VAF and sequencing depth at variant loci.

**Extended Data Figure 4.**
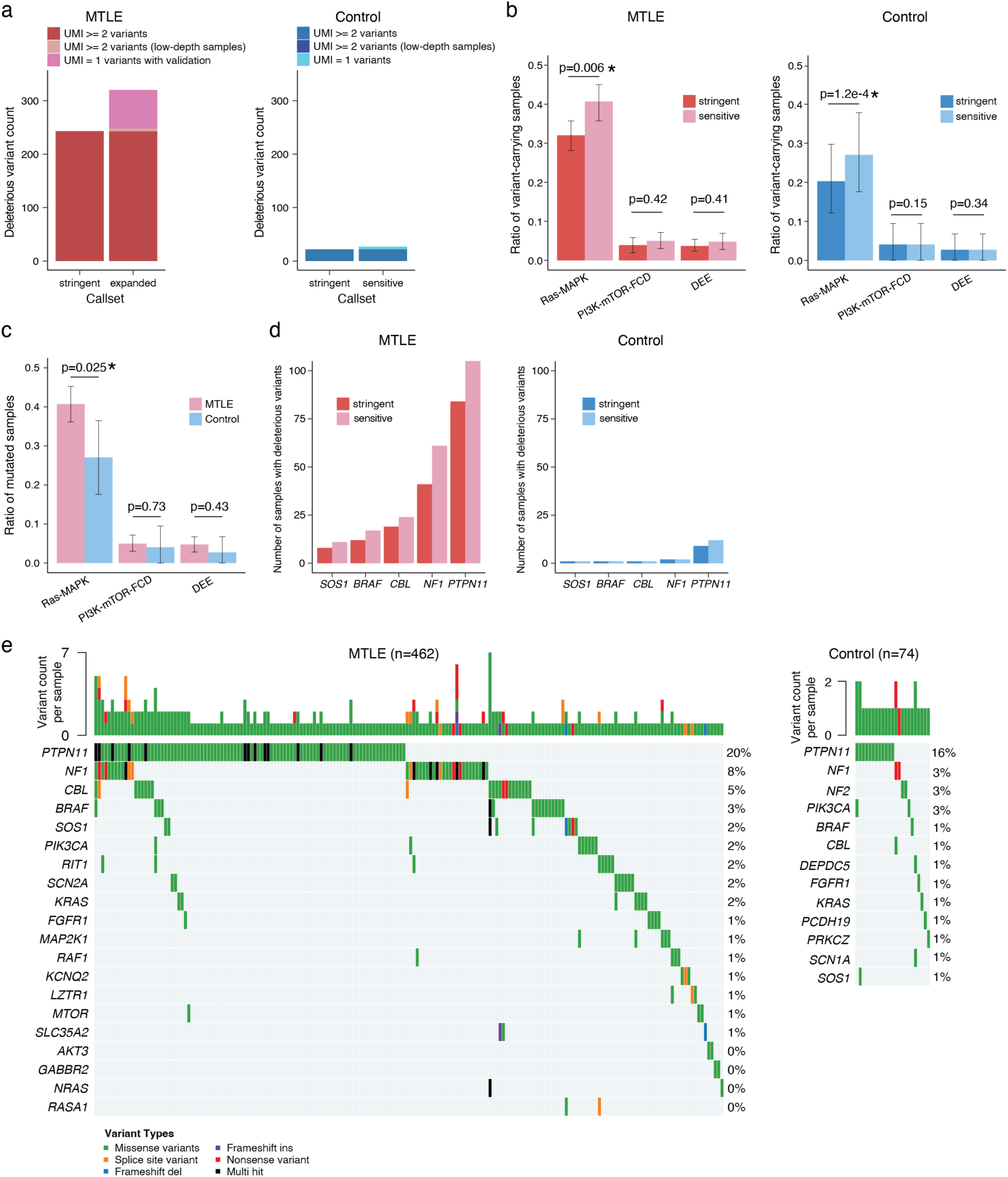
Increased deleterious somatic variant burden in the sensitive callset. (a) Distribution of deleterious somatic variants in the stringent and sensitive callsets. (b) Proportion of samples with deleterious somatic variants in each callset across gene groups (Two-sided Wilcoxon test; 95% CI via 1,000 bootstrap iterations). Asterisks indicate statistical significance. (c) Proportion of MTLE and control samples with deleterious somatic variants in the sensitive callset (Two-sided Wilcoxon test; 95% CI via 1,000 bootstrap iterations). Asterisks indicate statistical significance. (d) Number of samples with deleterious variants in in each callset for top mutated genes. (e) Oncoplot summarizing deleterious somatic variants in the sensitive callset. Top mutated genes are represented and variant types, variant burden per sample, and percentages of variants per gene are shown.

**Extended Data Figure 5.**
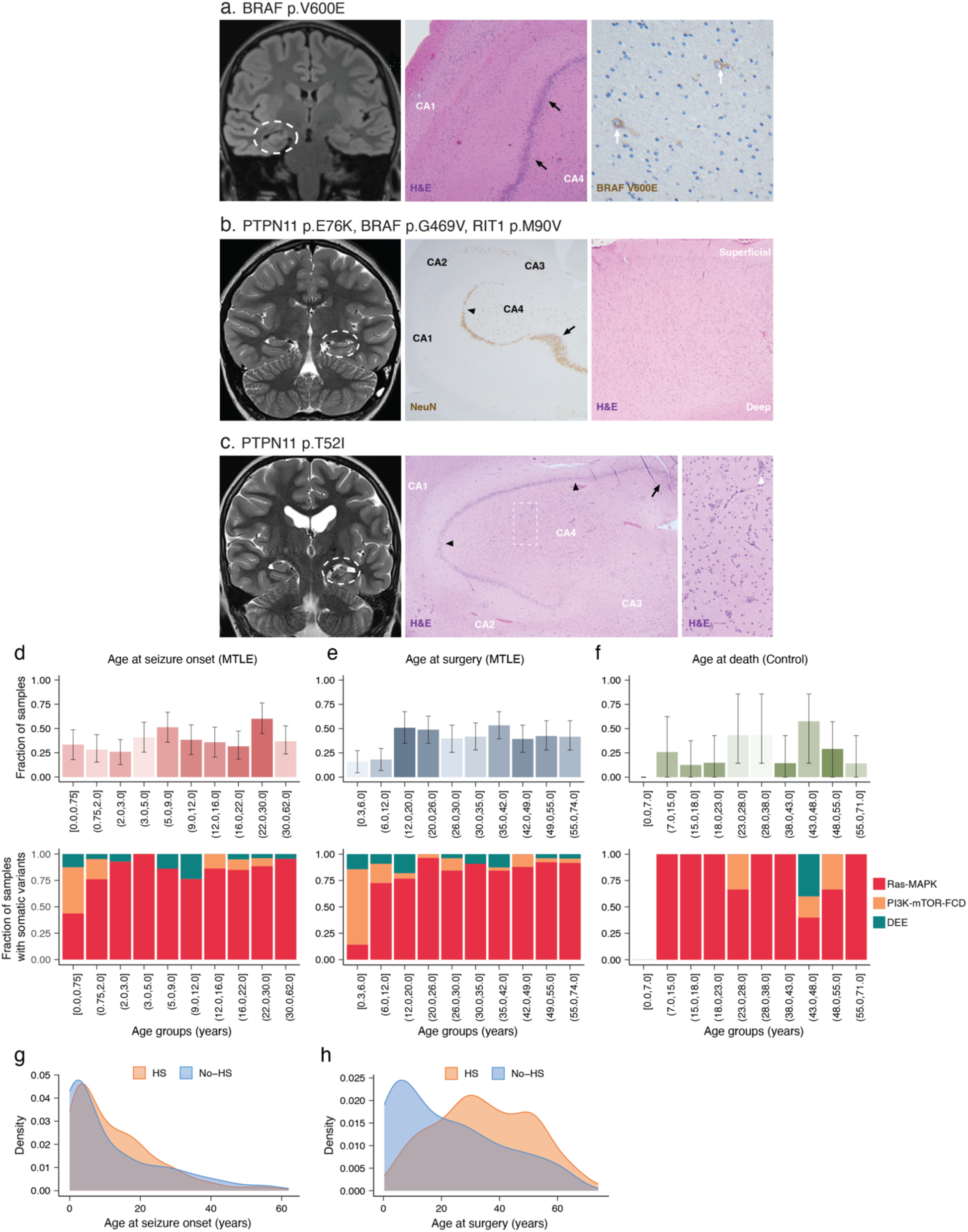
Association of deleterious somatic variants with age. (a-c) Representative coronal FLAIR MRI images and histopathological findings in three MTLE cases with deleterious Ras-MAPK variants. Dashed white circles highlight hippocampal volume loss and/or FLAIR hyperintensity on the presurgical MRI. Histopathologic hallmarks of HS including neuronal loss in the CA hippocampal subfields are present in all three cases. Black arrows and black arrowheads demarcate granule cell dispersion and loss in the dentate gyrus, respectively. (a) Neuronal loss in CA1 and granule cell dispersion (middle panel, 40X) and BRAF p.V600E immunoreactivity (white arrows) in neurons (right panel, 200X). (b) Widespread hippocampal neuronal loss and focal granule cell loss and dispersion (middle panel, 20X), and mild non-specific columnar arrangement of neurons in the adjacent temporal neocortex (right panel, 40X). (c) Widespread hippocampal neuronal loss and focal granule cell loss and dispersion (middle panel, 20X) and rare hypertrophic CA4 neurons (white arrowheads, 100X). Right panel is an amplification of the region in the dashed white box. (d-f) Proportions of samples with deleterious somatic variants stratified by deciles of age at seizure onset, age at time of surgery, and age at the time of death for controls (95% CI via 1,000 bootstrap iterations). Relative proportions of deleterious somatic variants in epilepsy-associated gene groups is shown in the bottom. (g-h) Incidence of HS (based on histopathology) across the spectrum for age at seizure onset (g) and age at the time of surgery (h).

**Extended Data Figure 6.**
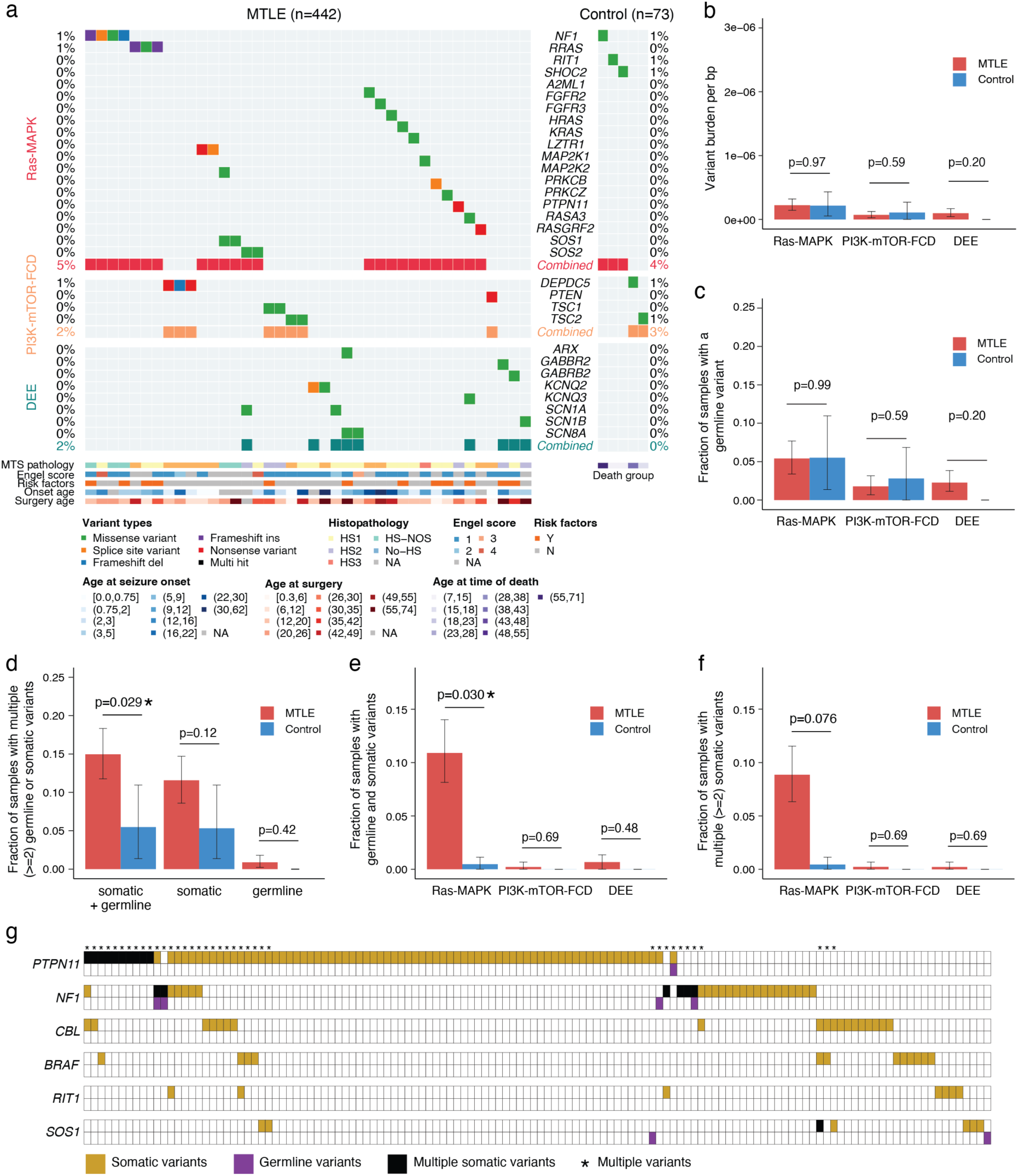
Distribution of deleterious germline variants. (a) Oncoplot summarizing deleterious germline variants detected in MTLE (n = 442) and control (n = 73) samples. Genes associated with Ras-MAPK signaling (top), PI3K-mTOR-FCD (middle), and DEE (bottom) are represented. The percentage of samples with deleterious germline variants is shown for MTLE (left) and controls (right). Variant types and clinical metadata are shown at the bottom. (b) Germline variant burden per base pair (bp) by gene group (Two-sided Wilcoxon test; 95% CI via 1,000 bootstrap iterations). (c) Proportion of samples with deleterious germline variants in each gene group (Two-sided Wilcoxon test; 95% CI via 1,000 bootstrap iterations). (d) Proportion of samples carrying multiple deleterious variants (Two-sided Wilcoxon test; 95% CI via 1,000 bootstrap iterations). Asterisks indicate statistical significance. (e) Proportion of samples with deleterious germline and somatic variants in the same gene group (Two-sided Wilcoxon test; 95% CI via 1,000 bootstrap iterations). Asterisks indicate statistical significance. (f) Proportion of samples with multiple deleterious somatic variants in the same gene group (Two-sided Wilcoxon test; 95% CI via 1,000 bootstrap iterations). (g) Oncoplot showing overlap of MTLE deleterious somatic and germline variants in highly mutated genes. Top and bottom rows for each gene show somatic and germline variants, respectively.

**Extended Data Figure 7.**
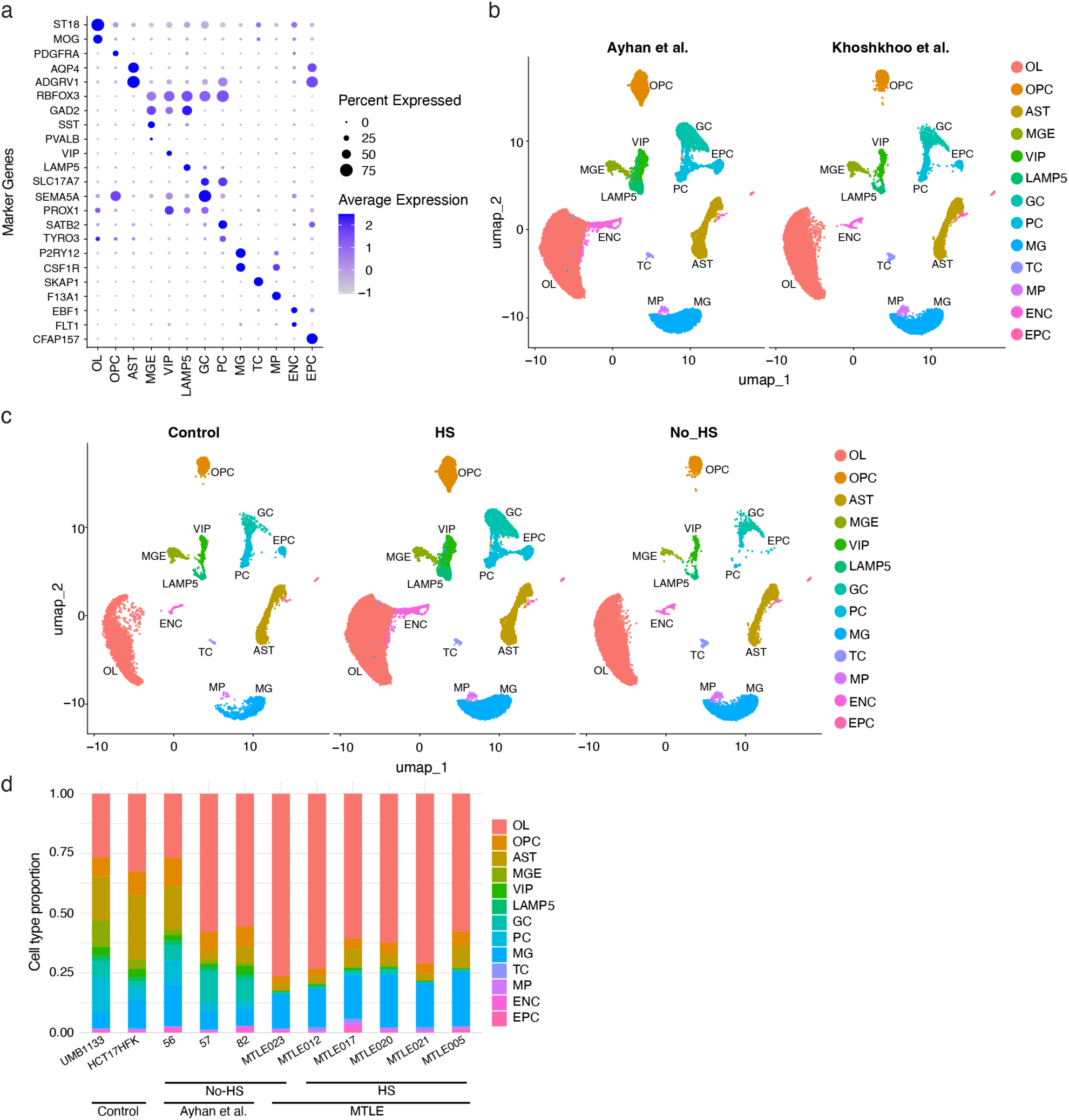
Distribution of snRNA-seq data by cohort and cell type. (a) Marker genes confirming snRNA-seq cluster identities. (b) Distribution of data generated for this paper and data previously published in Ayhan et al. (c) Distribution of data based on hippocampal histopathology. (d) Cell type proportions per sample.

**Extended Data Figure 8.**
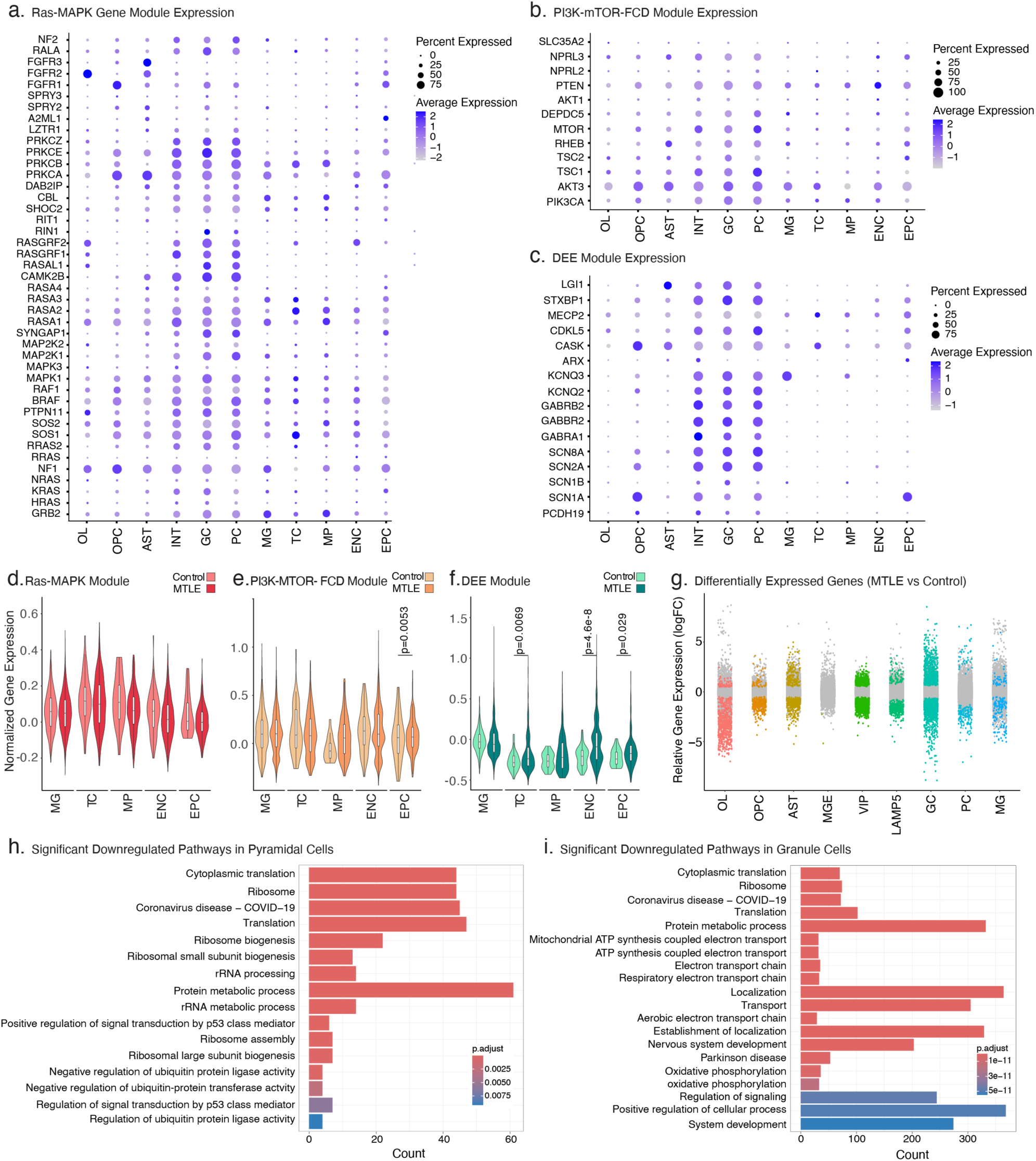
Differential expression of genes per cell type. (a-c) Mean expression of individual Ras-MAPK, PI3K-mTOR-FCD, and DEE genes included in the targeted sequencing panel per cell type. (d-f) Normalized expression levels for Ras-MAPK, PI3K-mTOR-FCD, and DEE genes in the targeted sequencing panel per cell type (Two-sided Wilcoxon test). P-values <0.05 are shown. (g) Relative expression of genes in MTLE samples compared to controls per cell type. Each dot represents one gene and colored dots represent differentially expressed genes in MTLE. (h-i) Top differentially downregulated gene ontology pathways in MTLE vs controls for pyramidal and granule cells. Only pathways with Benjamini-Hochberg-adjusted p-values <0.05 are shown.

**Extended Data Figure 9.**
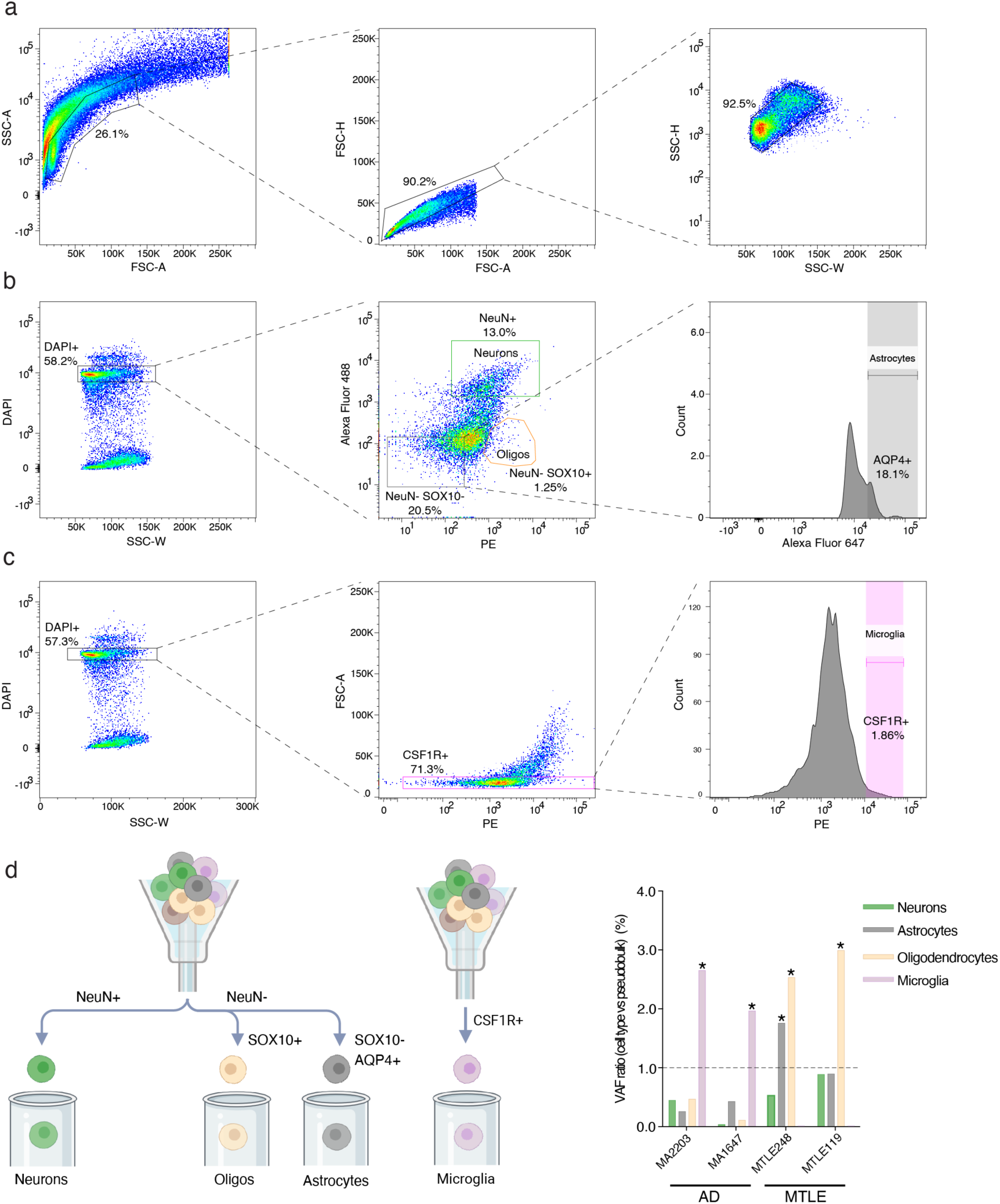
Nuclear sorting strategy and cell type enrichment in AD. (a) Gating strategy to remove debris and doublets. (b) Fluorescence Activated Nucleus Sorting (FANS) to isolate neurons, astrocytes, oligodendrocytes from brain tissue. (c) FANS to isolate microglia from brain tissue. (d) Relative abundance of somatic variants in each cell type in AD and MTLE brains as measured by ddPCR. Cell types with VAFs greater than the pseudobulk value (ratio of cell type-specific VAF over mean VAF for all cell types > 1, marked with asterisks) show relative enrichment in each sample.

## Supplementary Tables

**Supplementary Table 1. Sample and patient information.** Tissue source and basic demographic information for MTLE, neurotypical control, and AD brains used in duplex panel sequencing and snRNA-seq experiments are shown. For MTLE samples, important clinical variables including age at seizure onset and surgery, presence of key risk factors, radiological and histopathological classification of HS, and epilepsy surgery outcome are provided. For AD samples, Braak stage indicates degree of disease progression.

**Supplementary Table 2. Deleterious somatic and germline variant list.** List of deleterious somatic variants from the stringent and sensitive callsets and deleterious germline variants are shown. Variant type, predicted functional outcome, and annotations from dnSNP, GnomAD, ClinVar, EVE, REVEL, COSMIC, and SpliceAI are shown for every variant.

**Supplementary Table 3. Summary of orthogonal validation experiments.** Somatic variants experimentally tested for validation using amplicon sequencing and/or ddPCR are shown. The number of alternate (ALT) and reference (REF) reads and the calculated VAF for each amplicon and ddPCR experiment are indicated. Variants confirmed with >=2 amplicons or ddPCR are marked as validated (V), variants confirmed with only 1 amplicon are marked as likely validated (LV), and variants that are not confirmed experimentally are marked as not validated (NV).

**Supplementary Table 4. List of differentially expressed genes.** Genes with differential expression in MTLE vs control hippocampi, their relative expression in MTLE, and FDR-adjusted p-values are shown. Gene lists for different cell types are shown in different tabs.

**Supplementary Table 5. Summary of gene ontology analysis for differentially expressed genes.** Gene ontology terms for upregulated and downregulated genes in MTLE versus control hippocampi are shown. Gene ontology terms for different cell types and for upregulated and downregulated genes in MTLE are shown in different tabs.

**Supplementary Table 6. Brain regional enrichment analysis.** Abundance of MTLE variants in the hippocampus versus adjacent temporal neocortex was tested using ddPCR. Summary of the samples and variants tested and VAF in each brain region is shown.

